# Variation of anti-oomycete activity in *Pseudomonas* spp.: phenotypic characterization and comparative genomics

**DOI:** 10.64898/2026.04.24.720349

**Authors:** Ela Šarić, Anđela Miljanović, Petra Struški, Simone Oberhaensli, Livia Jerjen, Jurica Žučko, Heike Schmidt-Posthaus, Dora Pavić, Ivana Maguire, Julius Hermanns, Laure Weisskopf, Tobia Pretto, Ana Bielen

## Abstract

Pathogenic aquatic oomycetes *Aphanomyces astaci* and *Saprolegnia parasitica* represent a major threat to biodiversity and aquaculture production, but their interactions with host-associated microbes remain poorly understood. From a collection of bacterial isolates (*n* = 328) obtained from fish and crayfish hosts, we focused on *Pseudomonas* spp. (*n* = 66) and confirmed their previously reported strong inhibitory potential against *A. astaci* and *S. parasitica*. However, our results also revealed substantial inter-and intra-species variation in antagonism. To capture this variation, we selected eight isolates belonging to different *P. fluorescens*, *P. putida*, and *P. syringae* species groups and displaying contrasting levels of anti-oomycete activity for further phenotypic assays and comparative genomic analysis. Across these isolates, mycelial inhibition was markedly stronger against *A. astaci* than against *S. parasitica*, indicating species-specific differences in susceptibility. Comparative genomic analysis revealed substantial variation in biosynthetic gene cluster (BGC) repertoires among the analysed strains. Strongly inhibitory isolates carried candidate BGCs with similarity to characterised bioactive pathways, including pyoluteorin, rhizoxin, pyrrolnitrin, DAPG, and orfamide, alongside with multiple uncharacterised clusters that were either shared among inhibitory isolates or restricted to individual strains. All analysed genomes also contained clusters related to siderophore and HCN biosynthesis. However, *in vitro* assays showed that siderophore production was not clearly associated with inhibitory activity and that total inhibition was considerably higher than volatile-only inhibition, suggesting a substantial contribution of non-volatile, presumably diffusible, compounds. Altogether, our results suggest that *Pseudomonas* anti-oomycete activity is species-and strain-dependent and likely reflects different combinations of multiple metabolites, likely including diffusible compounds, rather than a single conserved mechanism. In conclusion, this study provides a foundation for future work aimed at resolving mechanisms underlying microbial antagonism toward aquatic oomycete pathogens.

## 1. Introduction

Disease dynamics depends on the complex interplay between the host immune system and the pathogen [1], but various environmental parameters can modulate disease development. Among them, environmental and host-associated microbes can play an important role in pathogenesis [2, 3]. During the invasion of the host, pathogens engage in complex and diverse interactions with other microbes, ranging from mutualism to competition/inhibition or hyperparasitism [4, 5]. Thus, microbe-microbe interactions can aid in the protection of the host against the pathogen, for example by competing with pathogens for resources or attachment sites, stimulating host immune defences, or producing antimicrobial compounds [6–8]. Mechanistic understanding of these interactions can explain patterns of disease resistance/susceptibility and open avenues for the development of novel disease control strategies.

Here, we have used *in vitro* co-cultivation assays and comparative genomics to study the antagonism between host-associated bacteria and oomycete pathogens of freshwater animals. Pathogenic oomycetes *Aphanomyces astaci* and *Saprolegnia parasitica* represent major threats to freshwater biodiversity and aquaculture [9–13]. *A. astaci*, the causative agent of crayfish plague, has decimated native European crayfish populations [14, 15]. *S. parasitica* causes saprolegniosis, a disease that affects all life stages of salmonids and other fish, resulting in high mortality and substantial economic losses in aquaculture, with losses in eggs and juvenile fish averaging around 10% and reaching up to 50% in severe outbreaks [16, 17].

Previous studies have shown that host-associated bacteria can inhibit pathogenic oomycetes *in vitro* [18–20] and *in vivo* [19, 21–23]. For example, *Aeromonas piscicola*, *A. sobria*, *Pantoea agglomerans*, and *Pseudomonas fluorescens* isolated from the skin of brown and rainbow trout infected by saprolegniosis showed high *in vitro* inhibition of *S. parasitica* [18]. Similarly, some bacteria from the cuticle of healthy crayfish individuals showed good inhibitory potential towards mycelial growth of *A. astaci*, and more than half of the identified inhibitors belonged to the genus *Pseudomonas* [20]. Further, *in vivo* studies showed that some *Pseudomonas* isolates can significantly reduce the mortality of salmon eggs caused by *S. diclina* [19] and can control *Saprolegnia* infection in adult rainbow trout [22].

Overall, in most of the studies focused on exploring the antagonistic potential of bacteria towards pathogenic oomycetes, *Pseudomonas* strains were highlighted as the most promising candidates [18–21]. However, most mechanistic insights into *Pseudomonas*-oomycetes interactions originate from plant-pathogenic oomycete systems [24, 25], where it was shown that pseudomonads can protect the host through (i) direct antagonism by secretion of anti-oomycete volatile or diffusible secondary metabolites [26]; (ii) occupation of ecological niches and competitive dominance, e.g., through iron binding by siderophores [6, 27]; and (iii) the activation of host defence responses [8].

Importantly, antagonistic activity can vary at the strain level, i.e., the observed inhibition sometimes differed markedly even among closely related *Pseudomonas* isolates [20, 28]. Such variation may reflect differences in genomic background and secondary metabolite repertoires, potentially shaped by processes such as gene gain, loss, or horizontal transfer. Comparative analysis of phylogenetically related strains with contrasting inhibitory phenotypes may therefore help identify candidate determinants of pathogen inhibition. Accordingly, the aims of this study were: (i) to quantify variation in the inhibitory activity of crayfish-and fish-associated *Pseudomonas* spp. against *A. astaci* and *S. parasitica*, and (ii) to use comparative genomics of *Pseudomonas* spp. with contrasting inhibitory phenotypes to *in silico* identify candidate biosynthetic gene clusters (BGCs) potentially underlying the observed inhibition. The overall study design is outlined in Figure 1.

**Figure 1.**
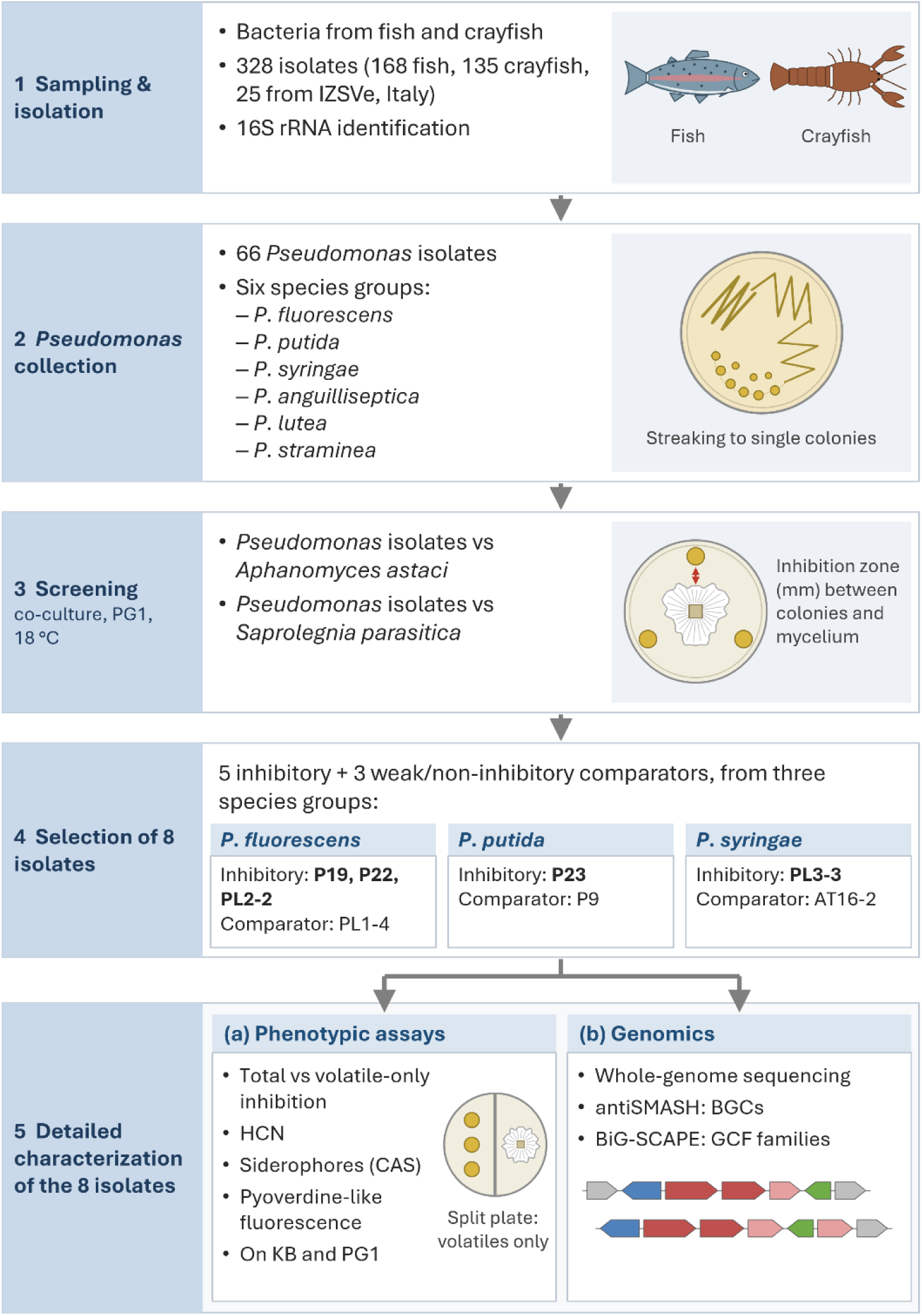
Study design. (1) Isolation of bacteria from fish and crayfish; (2) identification of *66 Pseudomonas* isolates; (3) co-culture screening against *A. astaci* and *S. parasitica*; (4) selection of five inhibitory and three comparator isolates; (5) phenotypic (a) and genomic (b) characterization of the selected isolates. The figure was prepared in Microsoft PowerPoint with the assistance of Claude Opus 5.5 (Anthropic), based on the study design and content provided by the authors.

## 2. Materials and methods

### 2.1. Collection of host-associated bacterial isolates

We have collected skin and exoskeleton biofilm samples of apparently healthy trout and crayfish, respectively, to culture host-associated bacterial isolates. No specific ethical approval for trout sampling was required because no fish were killed for the purposes of this study. Samples were collected from fish that had already been killed during routine commercial harvesting for human consumption at the fish farm. Crayfish were collected with permission obtained from the Ministry of Environmental Protection and Green Transition of the Republic of Croatia and in accordance with ethical standards.

A total of 168 and 135 bacterial isolates were obtained from the surface of fish and crayfish, respectively, and supplemented with 25 Pseudomonas isolates from various fish and crayfish organs obtained during routine diagnostic work at the Istituto Zooprofilattico Sperimentale delle Venezie (Legnaro, Italy). Sampling and culturing procedures, as well as the taxonomic composition of the collection, are described in Additional file 1. All isolates are listed in Additional file 2: Table S1. The collection served solely as a source of *Pseudomonas* isolates; isolates were therefore not compared according to sample origin or sampling procedure, host species or sampling date.

### 2.2. Preliminary identification of bacterial isolates by 16S rRNA gene sequencing

Bacterial isolates were preliminarily identified by sequencing of the nearly full-length 16S rRNA gene, followed by BLASTN comparison against the NCBI 16S rRNA database; details of DNA isolation, amplification and sequence analysis are provided in Additional file 1. For isolates assigned to the genus *Pseudomonas*, the closest BLASTN matches were used to assign isolates to species groups sensu Lalucat et al. [29], i.e. taxonomic units above the species level.

### 2.3. Plate-based co-culture inhibition assay

Anti-oomycete activity was assessed on solid PG1 medium [30] (per litre: 3 g peptone, 6 g glucose, 0.17 g MgCl_2_·x 6H_2_O, 0.145 g CaCl_2_·x 2H_2_O, 0.02 g FeCl_3_·x 6H_2_O, 0.37 g KCl, 0.055 g Na_2_EDTA and 6.7 mM sodium phosphate buffer, pH 7.0, solidified with 12 g agar) using Model 1 of the plate inhibition assay (Additional file 3: Figure S1) described by Orlić et al. [20], with some modifications. We have used *Aphanomyces astaci* Schikora, 1906 strain B, PsI genotype (isolate PEC 8), isolated from noble crayfish (*Astacus astacus*), Czech Republic (provided by F. Grandjean, University of Poitiers, Poitiers, France), and *Saprolegnia parasitica* Coker strain CBS (223.65) isolated from the northern pike (*Esox lucius*), Netherlands (provided by R. Galuppi, University from Bologna, Bologna, Italy). Bacterial isolates were pre-incubated alone at 22 °C for 48 h before the oomycete plug was added to the center of the plate. Co-cultures were then incubated at 18 ^°^C for nine days for *A. astaci* and four days for *S. parasitica*. Each assay was performed using three independent biological replicates per isolate. Endpoint inhibition zones were quantified in ImageJ software v1.53 [31] as the distance between the bacterial colony and the mycelial margin.

### 2.4. Split-plate assay to compare total (volatile and diffusible) and volatile-only inhibition

To compare total oomycete growth inhibition (mediated by both volatile and diffusible metabolites) with inhibition mediated by volatile metabolites only, *Pseudomonas* isolates and oomycetes were co-cultivated on solid PG1 medium [30] using two types of Petri dishes: split plates (allowing only the exchange of volatile compounds), and the regular non-split plates (allowing the exchange of both volatile and diffusible metabolites) (Additional file 3: Figure S2, [7]). Bacterial inocula were prepared by growing *Pseudomonas* isolates in liquid PG1 medium for 48 h at 28 °C and 150 rpm. Cultures were then standardized to an optical density at 600 nm (OD_600_) of 1 and 10 μL of each suspension was inoculated onto three positions per plate using sterile filter paper discs (d = 6 mm). Plates were incubated at 22 °C for 48 h. After the bacteria grew, mycelial plugs of *S. parasitica* or *A. astaci* were placed onto the other half of agar surface (Additional file 3: Figure S2). Plates were sealed with parafilm to prevent the loss of volatile compounds and incubated at 18 °C until the mycelium on the control split plates (without bacteria) reached the partition (two days for *S. parasitica* and five days for *A. astaci*); at that point, incubation was stopped on all plates, including the non-split plates, so that the mycelium did not come into contact with the bacterial colonies. All experiments were performed in three biological replicates for each isolate.

Mycelial growth was quantified by measuring colony area from scanned images using ImageJ v1.53 [31]. Total and volatile-mediated inhibition percentages were calculated relative to control plates of the same type (regular or split) inoculated with the oomycete only. Inhibition mediated by diffusible compounds was not quantified directly; its contribution was inferred from the difference between total and volatile-only inhibition. Inhibition percentages were calculated as follows:

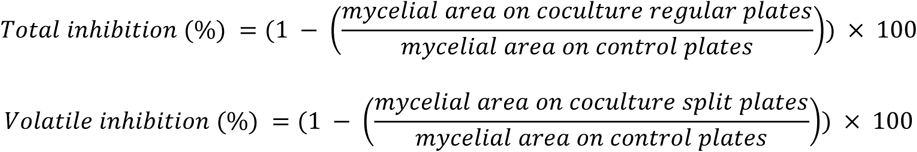

### 2.5. Detection of hydrogen cyanide (HCN) production

Hydrogen cyanide production was assessed qualitatively using HCN detection paper, which develops a blue colour in the presence of HCN. The detection reagent was prepared by dissolving copper(II) ethyl acetoacetate (50 g L⁻¹; Sigma Aldrich) and 4,4′-methylenebis(N,N-dimethylaniline) (50 g L⁻¹; Sigma-Aldrich) in chloroform (Fisher Chemicals). The solution was mixed until the compounds were fully dissolved, but for no longer than 15 min to minimise oxidation. The reagent was pipetted onto pre-cut Whatman filter-paper strips, which were dried overnight in a fume hood. The prepared detection papers were stored at 4 °C in a glass container protected from light with aluminium foil.

*Pseudomonas* isolates were precultured in LB broth, washed with sterile 0.9% NaCl, and adjusted to an OD₆₀₀ of 1.0. Aliquots (3 × 10 μL) were spotted onto King’s B (KB) and PG1 agar plates from which approximately half of the agar had been removed. HCN detection paper was placed in the agar-free half of the same Petri dish. Plates were sealed and incubated at 28 °C for 48 h, after which HCN production was evaluated visually based on blue discoloration of the detection paper. Sterile 0.9% NaCl served as the negative control. The experiment was performed in three biological replicates for each isolate and each medium.

### 2.6. Siderophore detection assays

#### 2.6.1. Qualitative and quantitative assessment of siderophore production

Siderophore production by *Pseudomonas* isolates was qualitatively assessed using the chrome azurol S (CAS) assay on solid KB and PG1 media [32–34]. *Pseudomonas* isolates were precultured in LB broth, washed with sterile 0.9% NaCl, and adjusted to an optical density of OD₆₀₀ = 1. Aliquots (1×10 μL) were spotted onto agar plates and incubated at 28 °C for 48 h. Siderophore production was indicated by a colour change of media from blue to yellow/orange. The experiment was performed in one biological replicate for each isolate and each medium.

Total siderophore activity was further quantified using a liquid CAS assay in 96-well microtiter plates [35]. Bacterial cultures were grown in 2 mL of liquid PG1 medium for 48 h at 28 °C and 150 rpm (i) without added iron (Fe–) (FeCl₃·x 6H₂O omitted from the standard PG1 medium formulation) and (ii) with supplementation of 0.02 g/L FeCl₃·x 6H₂O (Fe+, standard PG1 composition). After incubation, cultures were centrifuged (10 min, 4 °C, 4000 × g), and 100 μL of supernatant was mixed with 100 μL of CAS reagent (prepared according to [36], pH 6.8, filtered through a 0.2 μm membrane) in microtiter plates. Uninoculated Fe– and Fe+ PG1 medium variants were used as negative controls, and 30 mM EDTA served as a positive control. Absorbance at 630 nm was measured using a UV/Vis microplate reader (Molecular Devices, LLC 2023) with SoftMax Pro v7.2 software repeatedly over 2 h 30 min, with 5 s of shaking before each reading, until the signal stabilized; values at the plateau were used for calculations. Siderophore production was expressed as percentage of siderophore units using the formula:

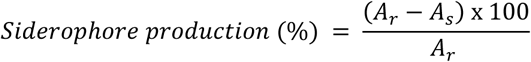

where *Ar* is the absorbance of the reference (uninoculated medium) and *As* is the absorbance of the sample. The number of biological replicates per isolate varied from two to seven, because additional experiments were performed for some isolates, whereas for others some replicates failed due to insufficient bacterial growth. Each biological replicate included two to four technical measurements.

#### 2.6.2. Qualitative and quantitative assessment of fluorescent metabolite production

The production of fluorescent metabolites, including pyoverdine, was assessed qualitatively on KB and PG1 agar media, while pyoverdine-associated fluorescence was quantified in liquid KB and PG1 media. *Pseudomonas* isolates were precultured in LB broth, washed with sterile 0.9% NaCl, and adjusted to an optical density of OD₆₀₀ = 1. Aliquots (10 μL) were spotted onto agar plates and incubated at 28 °C for 48 h. Colony fluorescence was documented under visible light and broad-spectrum UV illumination. As pyoverdine is the predominant fluorescent siderophore produced by fluorescent *Pseudomonas* spp. [37], the observed fluorescence may indicate pyoverdine production; however, other fluorescent metabolites may also contribute to the detected signal. The experiment was performed in one biological replicate for each isolate and each medium.

For quantitative analysis, aliquots of 5 μL bacterial suspension were inoculated into 195 μL of sterile medium in 96-well microplates and incubated for 48 h at 28 °C with shaking (110 rpm). Following incubation, optical density at 600 nm and fluorescence at wavelengths characteristic of pyoverdine (excitation 405 nm/emission 460 nm) were measured using a Cytation 5 microplate reader (BioTek Instruments, USA). Both OD and fluorescence values were corrected by subtracting the corresponding blank values. Pyoverdine-associated fluorescence was calculated for each well by dividing blank-corrected fluorescence by blank-corrected OD₆₀₀ and was expressed as relative fluorescence units per unit of OD₆₀₀ (RFU/OD₆₀₀). The experiment was performed in three biological replicates for each isolate and each medium.

### 2.7. Statistical analysis

Statistical analyses were performed in Python 3.12.13 using Google Colab and the pandas (v2.2.2), NumPy (v2.0.2), SciPy (v1.16.3), scikit-posthocs (v0.13.0), and openpyxl (v3.1.5) packages. Biological replicates were used as the unit of statistical analysis. Where technical replicates were performed (liquid CAS assay), their values were averaged for each biological replicate. Because the number of biological replicates was small and the assumptions required for parametric tests could not be reliably established, non-parametric tests were used. Siderophore production under iron-limited (Fe−) and iron-supplemented (Fe+) conditions was compared using the paired Wilcoxon signed-rank test. Differences among bacterial isolates were assessed separately within each iron condition using the Kruskal–Wallis test. Significant Kruskal–Wallis results were followed by pairwise Dunn’s tests with Holm adjustment for multiple comparisons. General antagonistic activity, total and volatile-only inhibition, and pyoverdine-associated fluorescence were analysed using the same non-parametric approach. Statistical significance was set at *p* < 0.05.

### 2.8. Genome sequencing and bioinformatic analyses

The eight selected *Pseudomonas* isolates were revived from glycerol stocks and streaked on PG1 plates to obtain individual colonies. A single colony of each isolate was then inoculated into 2 mL of liquid PG1 medium and incubated at 21 °C with shaking for two days. Pellets containing approximately 10⁹ cells each (corresponding to 1 mL of *Pseudomonas* culture of OD₆₀₀ = 1) were prepared for each isolate and used for total genomic DNA extraction with Qiagen MagAttract HMW DNA Kit (Qiagen, 67563) following the guidelines for Gram Negative bacteria (Qiagen User Guide, HB-1523-006_HB_MagAttract_HMW_DNA_03/20) and a Thermo Fisher Scientific Kingfisher Apex robot. The resulting bacterial genomic DNA was assessed for quantity, quality and purity using a Qubit 4.0 flurometer (Qubit dsDNA HS Assay kit; Q32851, Thermo Fisher Scientific), an Advanced Analytical FEMTO Pulse instrument (Genomic DNA 165 kb Kit; FP-1002-0275, Agilent) and a Denovix DS-11 UV-Vis spectrophotometer, respectively.

PacBio HiFi library preparation and Revio sequencing were performed at the Next Generation Sequencing Platform, University of Bern, following the general workflow described by Jordi et al. [38]. Libraries were prepared according to the HiFi plex prep kit 96 protocol 103-418-800 REV02, and DNA shearing followed PacBio technical note 102-326-575 REV01. Sequencing complexes were prepared in SMRT Link v25 using the Revio SPRQ polymerase kit (PN 103-496-900) and clean-up beads (PN 102-158-300) and loaded onto a Revio SPRQ sequencing plate (PN 102-326-552) with SMRT Cell 25M trays (PN 102-202-200). Sequencing was performed with a 24 h movie time.

Demultiplexed HiFi reads in fastq format were assembled using Flye 2.9.5 [39] with default parameters for hifi data (--pacbio-hifi). Assemblies were annotated with bakta v1.9.4 using database v5.1 “full” [40], circular chromosomes were rotated so that they start with the dnaA gene using a custom python script. The genome assemblies of the eight selected *Pseudomonas* strains generated in this study have been deposited in DDBJ/ENA/GenBank under BioProject accession number PRJNA1494142. The accession numbers of the genome assembly versions described in this paper are provided in Table 1.

**Table 1.** Overview of genome characteristics of selected *Pseudomonas* strains. Species names were assigned through NCBI taxonomy upon deposition of the genome assemblies in GenBank; they were not determined in this study, in which isolates are referred to by species group. PL1-4 was assigned to the *P. fluorescens* group based on its 16S rRNA gene sequence, but whole-genome phylogeny placed it outside the subclade comprising the other *P. fluorescens*-group isolates (Additional file 3: Figure S9); it is therefore referred to as *Pseudomonas* sp. in the text.

| Isolate | **P19** | **P22** | **PL2-2** | **PL1-4** | **P23** | **P9** | **PL3-3** | **AT16-2** |
|---|---|---|---|---|---|---|---|---|
| **Accession number** | JCFXCP010000000 | JCFXCS010000000 | JCFXCN010000000 | JCFXCU010000000 | JCBPDU010000000 | JCESCB010000000 | JCFXCQ010000000 | JCFXCR010000000 |
| **Species group** | *P. fluorescens* | *P. fluorescens* | *P. fluorescens* | *Pseudomonas* sp. | *P. putida* | *P. putida* | *P. syringae* | *P. syringae* |
| **Species** | *P. idahonensis* | *P. paracarnis* | *P. chlororaphis* | *P. fluorescens* | *P. wadenswilerensis* | *P. mosselii* | *P. fragariae* | *P. congelans* |
| **Total assembly size (bp)** | 6,871,141 | 6,728,914 | 6,939,645 | 6,486,735 | 6,030,240 | 6,102,432 | 6,229,185 | 5,709,200 |
| **Number of contigs** | 1 | 3 | 1 | 2 | 1 | 3 | 1 | 1 |
| **Assembly structure** | circular chromosome | 3 contigs | circular chromosome | circular chromosome + circular plasmid | circular chromosome | 3 contigs | circular chromosome | circular chromosome |
| **GC content (%)** | 62.2 | 59.6 | 62.6 | 60.8 | 62.3 | 64.2 | 59.0 | 59.4 |
| **CDS** | 6,037 | 6,045 | 6,099 | 5,753 | 5,427 | 5,269 | 5,338 | 4,888 |
| **rRNA genes** | 16 | 19 | 16 | 19 | 19 | 22 | 16 | 16 |
| **tRNA genes** | 73 | 73 | 68 | 73 | 74 | 79 | 63 | 63 |
| **Number of BGCs** | 23 | 18 | 17 | 11 | 14 | 15 | 15 | 10 |
| **Contig length (bp)/average coverage** | 6,871,141 (37x) | 5,384,563 (90x)<br>1,296,698 (83x)<br>47,653 (173x) | 6,939,645 (32x) | 6,319,495 (345x)<br>167,240 (96x) | 6,030,240 (82x) | 5,896,008 (74x)<br>172,713 (89x)<br>33,711 (285x) | 6,229,185 (82x) | 5,709,200 (190x) |

Phylogenetic placement of the sequenced genomes was performed using the “Insert Set of Genomes Into SpeciesTree” (v2.2.0) app within the KBase platform (Knowledge Basel, https://www.kbase.us/) platform [41]. The phylogenetic tree was generated based on a set of 49 universal core genes (Clusters of Orthologous Groups, or COGs) defined by the KBase reference database [42]. Hundred most similar reference genomes from the KBase/RefSeq database were included in the tree (**“**Neighbor Public Genome Count” = 100). Curated alignment was trimmed using Gblocks and concatenated [43]. A maximum-likelihood phylogenetic tree was inferred from the concatenated alignment using FastTree2 and the fastest settings [44].

The obtained genomes were further analysed using antiSMASH for bacteria (version 8.0.4; [45]), with default settings and comparison to the MIBiG database enabled. The detected clusters were subsequently compared within and between genomes using BiG-SCAPE (version 2.0.2; [46]), i.e. the identified BGCs were grouped into gene cluster families (GCFs) based on domain architecture and sequence similarity, at a category-level pairwise distance cutoff of 0.3.

In addition, the genomic neighborhood of the identified BGCs was analysed in D3GB (https://d3gb.usal.es/; [47]) by examining five genes upstream and downstream of each cluster for nearby mobility-related genes, including transposases, IS element proteins, integrases, retrotransposon proteins, and other mobile element-associated proteins.

## 3. Results

### 3.1. Inhibition of *A. astaci* and *S. parasitica* mycelial growth by host-associated *Pseudomonas* isolates

Based on the reported strong anti-oomycete activity of *Pseudomonas* spp. [18–21], we performed inhibition assays on the representatives of this genus (Additional file 4: Table S2). From the total collection of 168 fish-and 135 crayfish-derived bacterial isolates, we selected all *Pseudomonas* strains (4 from fish and 37 from crayfish). These were supplemented with 25 additional *Pseudomonas* isolates obtained from fish (23), crayfish (1) and aquarium water (1) during routine diagnostics at Istituto Zooprofilattico Sperimentale delle Venezie, Venice, Italy. This resulted in a final collection of 66 *Pseudomonas* isolates tested for anti-oomycete activity.

The isolates in our collection were assigned to six species groups following the classification of Lalucat et al. [29]: *P. fluorescens* (39.4%), *P. putida* (36.4%), *P. syringae* (9.1%), *P. anguilliseptica* (6.1%), *P. lutea* (4.5%) and *P. straminea* (3%). One isolate (1.5%) could not be assigned to any of these groups and was designated as not determined (ND).

The highest number of isolates displaying strong inhibition of oomycete mycelial growth, particularly against *A. astaci*, was found within the *P. fluorescens* group, followed by the *P. putida* and *P. syringae* groups (Figure 2, Additional file 4: Table S2). Importantly, inhibitory activity varied substantially within individual species groups. Isolates belonging to the same *Pseudomonas* species group often exhibited pronounced differences in their inhibitory potential against freshwater pathogenic oomycetes (Figure). On the other hand, inhibition of *A. astaci* was stronger than that of *S. parasitica* across the tested *Pseudomonas* isolates (Figure 2). The strongest inhibitors of *A. astaci* were P19 (*P. fluorescens* group) and P23 (*P. putida* group), whereas PL2-2 (*P. fluorescens* group) and PL3-3 (*P. syringae* group) showed the strongest inhibition of *S. parasitica*.

**Figure 2.**
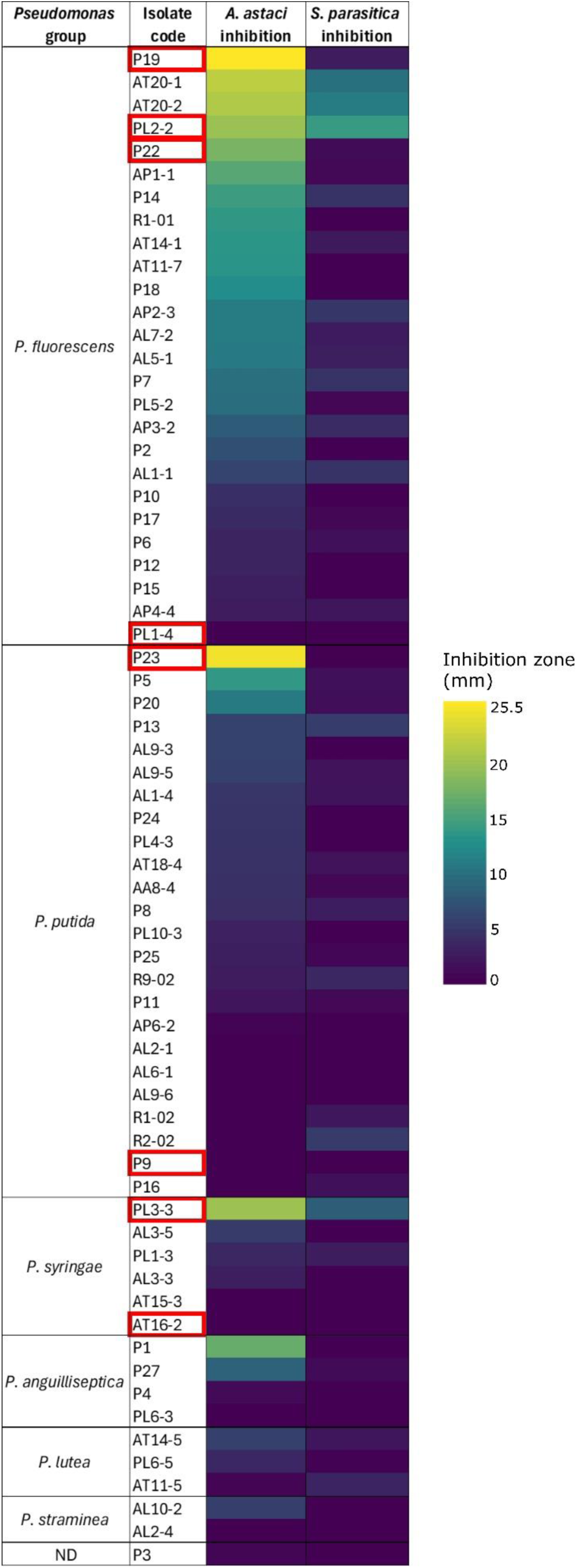
Heatmap showing the inhibitory activity of 66 *Pseudomonas* isolates against the mycelial growth of *A. astaci* and *S. parasitica*. Isolates are grouped according to their assignment to *Pseudomonas* species groups based on 16S rRNA gene sequences, following Lalucat et al. [29]. Isolate P3 could not be assigned to any of the previously defined groups and is therefore marked as ND. Each row represents a single isolate, and colour intensity reflects inhibition zone width (mm), ranging from low (dark purple) to high (yellow). The eight isolates selected for further phenotypic and genomic characterization are outlined in red.

Based on the screening results, eight *Pseudomonas* isolates were selected for further phenotypic and genomic characterization according to the following criteria: (i) the strongest inhibitors in the screening assay, including both isolates that inhibited both oomycetes (PL2-2 and PL3-3) and isolates that inhibited *A. astaci* only (P19, P22 and P23); (ii) representation of the three species groups that contained strong inhibitors (*P. fluorescens*, *P. putida* and *P. syringae*); and (iii) for each of these groups, one phylogenetically related isolate with weak or no inhibitory activity, included as a comparator (PL1-4, P9 and AT16-2). The selected isolates are outlined in red in the screening heatmap (Figure 2). No pronounced inhibitors were found in the *P. anguilliseptica*, *P. lutea* and *P. straminea* groups (Figure 2), the latter two comprising only three and two isolates, respectively, so phylogenetically matched inhibitor/non-inhibitor pairs could not be formed within them. Further analyses were therefore restricted to the *P. fluorescens* (inhibitors P19, P22 and PL2-2; weak/non-inhibitor PL1-4, hereafter referred to as *Pseudomonas* sp. PL1-4 because whole-genome phylogeny did not confirm its affiliation with this group; see Section 3.2.2), *P. putida* (inhibitor P23; weak/non-inhibitor P9) and *P. syringae* (inhibitor PL3-3; weak/non-inhibitor AT16-2) groups.

The inhibitory activity of the eight selected isolates, i.e. the same screening data as shown in Figure 2, is presented in more detail in Figure 3, including individual replicates and statistical comparisons. Significant differences among isolates were detected for both *A. astaci* (Kruskal–Wallis, *H* = 27.03, *p* < 0.001) and *S. parasitica* (*H* = 21.02, *p* = 0.0037). Dunn’s post hoc test revealed significantly greater inhibition of *A. astaci* by *P. fluorescens* P19 compared with *P. syringae* AT16-2, *P. putida* P9, and *Pseudomonas* sp. PL1-4 (Holm-adjusted *p* = 0.024 for all comparisons). For *S. parasitica*, no pairwise differences remained significant after Holm correction.

**Figure 3.**
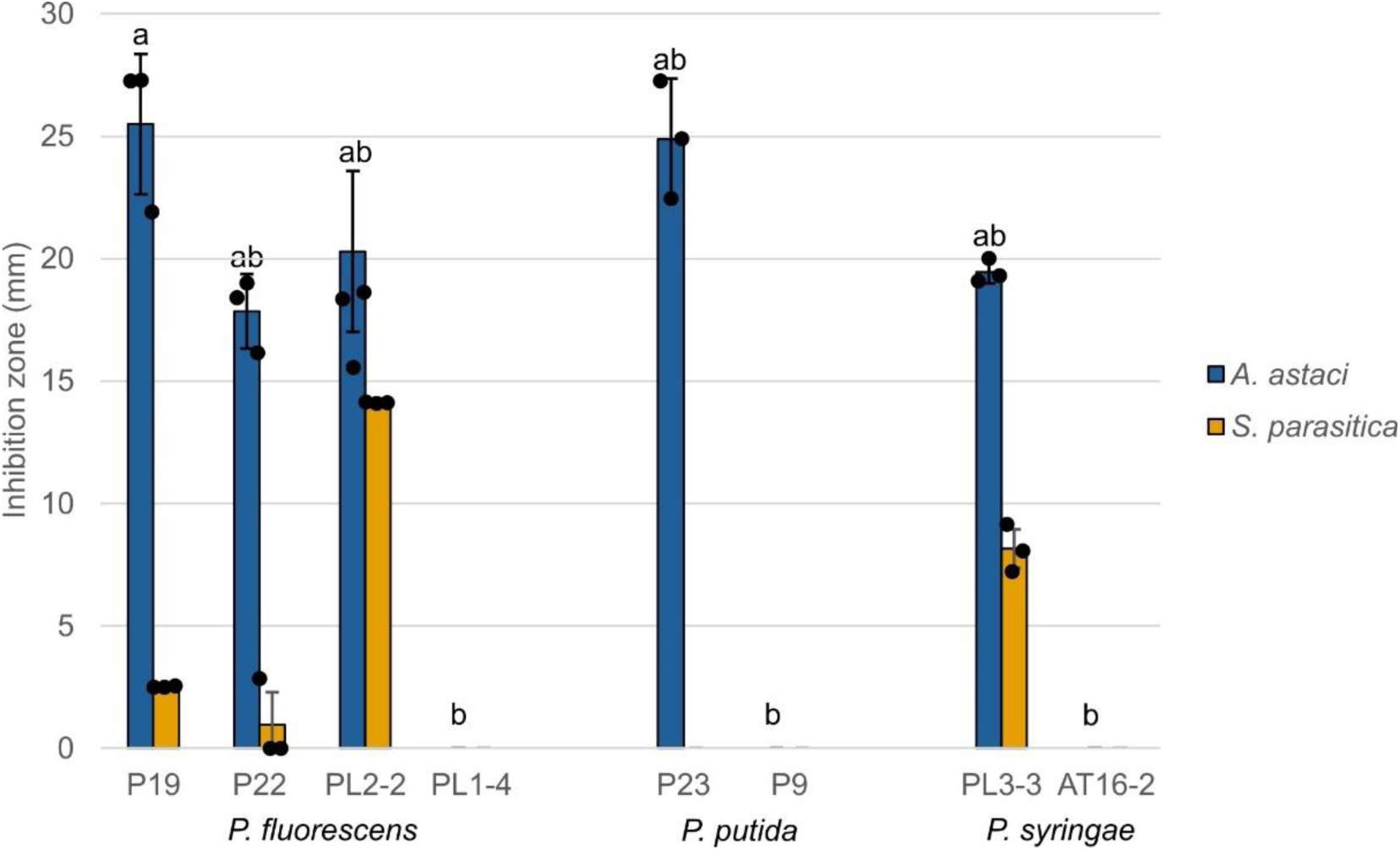
Effect of *Pseudomonas* strains on mycelial growth of *A. astaci* and *S. parasitica* analysed by plate-based co-culture inhibition assay. Values represent mean ± standard deviation (*n* = 3); individual measurements are shown as black dots. Letters above the *A. astaci* bars indicate the results of Dunn’s post hoc test with Holm adjustment following a significant Kruskal–Wallis test (*p* < 0.05); isolates sharing a letter do not differ significantly. For *S. parasitica*, no pairwise differences remained significant after Holm adjustment.

### 3.2. Phenotypic and genomic characterization of selected *Pseudomonas* isolates with contrasting inhibitory phenotypes

#### 3.2.1. Phenotypic components of inhibition

To further investigate the observed antagonism, we compared total and volatile-only inhibition and analysed HCN and siderophore production and fluorescent metabolites, including pyoverdine-associated fluorescence, by the selected *Pseudomonas* strains (Figure 4).

**Figure 4.**
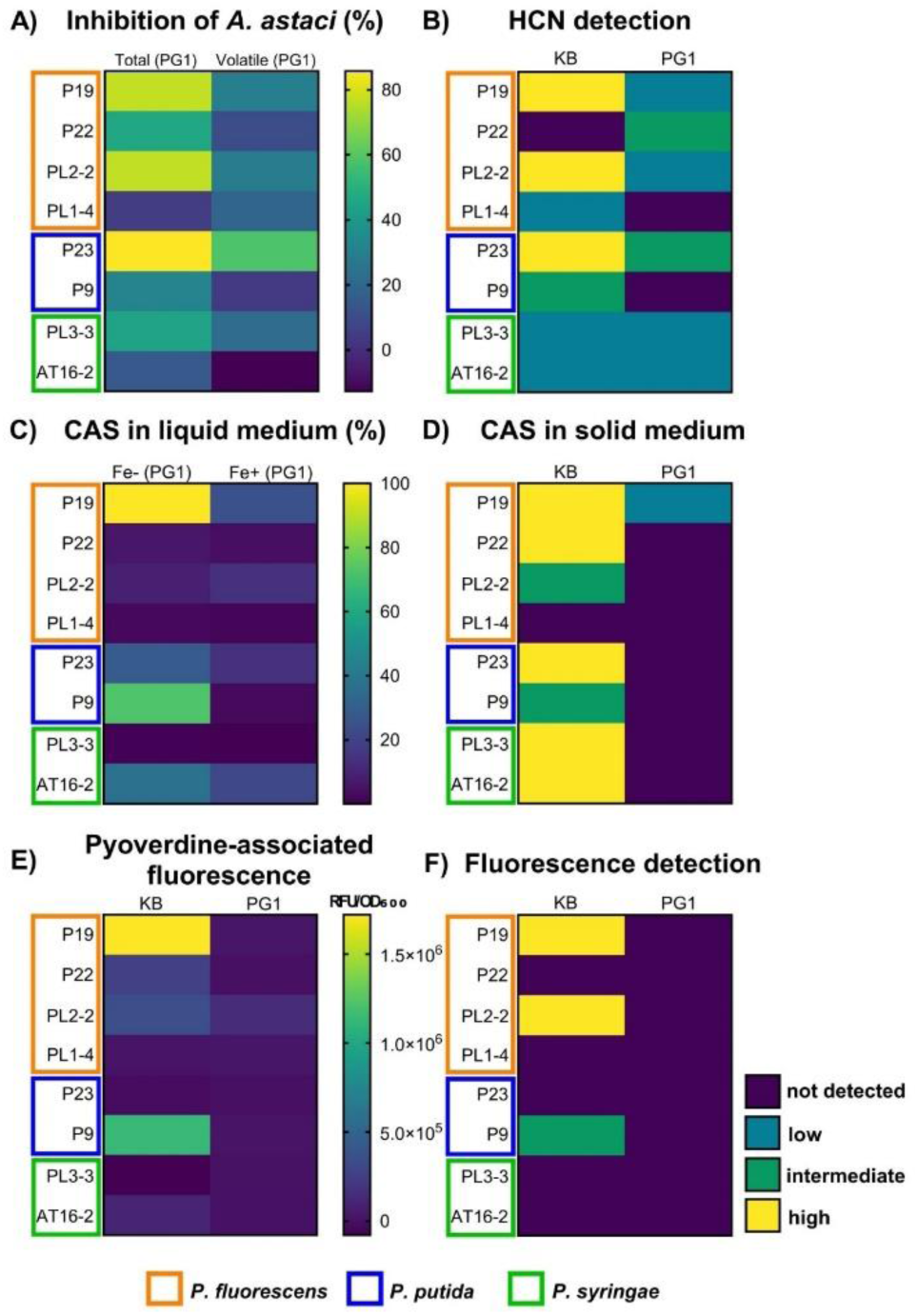
Integrated phenotypic characterization of the eight selected *Pseudomonas* isolates. The heatmap summarizes the results of quantitative (A, C, E) and qualitative (B, D, F) assays. (A) Total and volatile-only inhibition of *A. astaci* (%) on PG1 medium. (B) HCN production on KB and PG1 media. (C) Siderophore activity determined using the quantitative CAS assay in liquid PG1 medium under iron-limited (Fe−) and iron-supplemented (Fe+) conditions, expressed as percentages of siderophore units relative to uninoculated PG1 medium of the same composition (%). (D) Qualitative siderophore detection using the CAS assay on solid KB and PG1 media. (E) Pyoverdine-associated fluorescence measured in liquid KB and PG1 media, expressed as relative fluorescence units (RFU) normalized to OD₆₀₀. (F) Qualitative fluorescence detection on solid KB and PG1 media. Color scales represent the measured values in quantitative assays and visually assessed reaction intensities in qualitative assays. Isolates are affiliated to species groups.

The results of the split-plate assay have shown that *Pseudomonas* isolates generally inhibited the mycelial growth of *A. astaci* more strongly than that of *S. parasitica* (Additional file 3: Figure S3, Additional file 5: Table S3), consistent with the results of plate-based co-culture inhibition assay (Figure 3). Figure 4A presents the split-plate results for *A. astaci* only, as a heatmap, whereas detailed results are shown in Additional file 3: Figure S3A. Significant differences among isolates were detected for both total inhibition (Kruskal–Wallis, *H* = 19.95, *p* = 0.0057) and volatile-only inhibition (*H* = 15.01, *p* = 0.0359) of *A. astaci*. For *P. fluorescens* P19 and PL2-2, total inhibition (76.8% and 76.5%, respectively) was considerably higher than volatile-only inhibition (28.4% and 28.06%, respectively), whereas *P. putida* P23 showed substantial volatile-only inhibition (58.2% compared with 85.8% of total inhibition). For *P. syringae* PL3-3, total inhibition reached 44.1%, whereas volatile-only inhibition was 21.01%. Dunn’s post hoc test revealed significantly greater total inhibition by *P. putida* P23 compared with *Pseudomonas* sp. PL1-4 (Holm-adjusted *p* = 0.0339), whereas no pairwise differences in volatile-only inhibition remained significant after Holm correction. For *S. parasitica* (Additional file 3: Figure S3B), no significant differences among isolates were detected in total inhibition (Kruskal–Wallis, *H* = 13.99, *p* = 0.0514), whereas volatile-mediated inhibition differed significantly among isolates (*H* = 16.37, *p* = 0.0219). However, Dunn’s post hoc test revealed no significant pairwise differences after Holm correction.

To further characterize the volatile component of antagonism, HCN production by *Pseudomonas* strains was assessed on KB and PG1 media (Figure 4B, Additional file 3: Figure S4). HCN production differed between the two media, with stronger positive reactions generally observed on KB than on PG1 (Figure 4B). Strong positive HCN reactions were detected in most isolates exhibiting a substantial contribution of volatile metabolites to the overall inhibition, including *P. fluorescens* P19 and PL2-2, as well as *P. putida* P23, whereas isolates with little volatile-mediated inhibition generally showed weak or no detectable HCN production.

Next, siderophore production by the selected isolates was analysed both qualitatively and quantitatively. It was first qualitatively assessed using the CAS assay on solid KB and PG1 media. On KB medium, seven of the eight isolates showed a positive CAS reaction, with the strongest responses observed for *P. fluorescens* P19 and P22, *P. putida* P23, and *P. syringae* PL3-3 and AT16-2 (Figure 4D, Additional file 3: Figure S5). In contrast, on PG1 medium, a weak positive reaction was detected only for *P. fluorescens* P19. No CAS reaction was detected for *Pseudomonas sp.* PL1-4 on either medium.

Siderophore activity was further quantified using the liquid CAS assay in iron-limited (Fe−) and iron-supplemented (Fe+) PG1 media (Figure 4C, Additional file 3: Figure S6, Additional file 5: Table S3).Overall, siderophore production was significantly higher under iron-limited conditions (paired Wilcoxon signed-rank test, *W* = 32, *p* = 0.0273). Under both Fe− conditions, siderophore production differed significantly among isolates (Kruskal–Wallis, *H* = 23.27, *p* = 0.00153), with isolate *P. fluorescens* P19 producing significantly more siderophores than *Pseudomonas* sp. PL1-4 and *P. syringae* PL3-3 (Dunn’s test, Holm-adjusted *p* = 0.034 and 0.017, respectively). Under Fe+ conditions, siderophore production also differed among isolates (Kruskal–Wallis, *H* = 26.13, *p* = 0.00048), with a significant difference detected between *P. fluorescens* P22 and *P. putida* P9 (Dunn’s test, Holm-adjusted *p* = 0.039). However, siderophore production varied considerably among isolates and showed no clear relationship with either species group or previously determined anti-oomycete activity (Figure 4C).

In addition to the CAS assay, broad-range fluorescence was assessed on solid KB and PG1 media as a qualitative phenotypic screen for fluorescent siderophore pyoverdine and other fluorescent metabolites. Strong fluorescence was observed only for *P. fluorescens* isolates P19 and PL2-2 on KB media, whereas isolate *P. putida* P9 exhibited intermediate fluorescence (Figure 4E, Additional file 3: Figure S7). No fluorescence was detected on PG1 media. Quantification of pyoverdine-associated fluorescence in liquid cultures revealed significant differences among isolates on both KB (Kruskal–Wallis, *H* = 21.81, *p* = 0.0027) and PG1 media (*H* = 20.56, *p* = 0.0045) (Figure 4E, Additional file 3: Figure S8, Additional file 5: Table S3). On KB medium, *P. fluorescens* P19 exhibited the highest fluorescence intensity, followed by *P. putida* P9. Dunn’s post hoc analysis identified significant differences only between *P. fluorescens* P19 and *P. syringae* PL3-3 (*p* = 0.0077) and between *P. putida* P9 and *P. syringae* PL3-3 (*p* = 0.0492). On PG1 medium, *P. fluorescens* PL2-2 displayed the highest fluorescence intensity and differed significantly from *P. fluorescens* P22 (*p* = 0.0404) and *P. putida* P23 (*p* = 0.0280). The remaining isolates exhibited lower or undetectable fluorescence on both media.

Overall, the presented results (Figure 4) revealed considerable variation in inhibitory activity and metabolite-associated phenotypes among the eight selected *Pseudomonas* isolates. For most isolates, total inhibition of *A. astaci* exceeded volatile-only inhibition, suggesting substantial contribution of diffusible compounds to antagonistic phenotype. HCN and siderophore production was generally more pronounced on KB than on PG1, demonstrating that these phenotypes depend on nutrient availability. Strongly inhibitory *P. fluorescens* P19 stood out in almost all assays. Notably, *P. putida* P9 exhibited detectable fluorescence despite its relatively low inhibitory activity, while *Pseudomonas* sp. PL1-4 and *P. syringae* AT16-2 generally showed weak or undetectable responses across most assays. Nevertheless, no single assessed trait was consistently associated with anti-oomycete activity.

#### 3.2.2. Comparative genomic analysis

High-quality genome assemblies of the eight selected *Pseudomonas* strains are summarised in Table 1. Genome sizes ranged from 5,709,200 bp in *P. syringae* AT16-2 to 6,939,645 bp in *P. fluorescens* PL2-2. Six strains could be assembled into a circular chromosome, or a circular chromosome and a plasmid (PL1-4) (Table 1). The assemblies of *P. fluorescens* P22 and *P. putida* P9 comprised three contigs, but it could not be reliably determined whether all contigs were of chromosomal origin or whether some were plasmid-derived. The GC content ranged from 59.0% to 64.2%, and the number of predicted coding sequences (CDS) ranged from 4,888 to 6,099. Ribosomal RNA gene counts varied between 16 and 22, and tRNA gene counts between 63 and 79. These values are consistent with previously reported genome sizes and GC contents for members of the *P. fluorescens, P. putida* and *P. syringae* groups [48–50]. Whole-genome phylogenetic analysis of the strains was largely consistent with the initial 16S rRNA-based group assignment for seven of the eight isolates, which grouped within the *P. fluorescens*, *P. syringae*, and *P. putida* clades (Additional file 3: Figure S9). The only exception was the isolate PL1-4, which formed a separate branch outside the subclade comprising the inhibitory *P. fluorescens*-group isolates P19, P22 and PL2-2, together with *Pseudomonas* sp. StFLB209, originally isolated from the potato phyllosphere [51]. Because its group affiliation could not be resolved from the genome-based phylogeny, PL1-4 is referred to as *Pseudomonas* sp. throughout this study, although it was assigned to *the P. fluorescens* group based on its 16S rRNA gene sequence and to *P. fluorescens* by NCBI taxonomy upon genome deposition (Table 1). Differences between PL1-4 and the inhibitory *P. fluorescens* isolates may therefore also reflect this greater phylogenetic distance in addition to differences in inhibitory phenotype. By contrast, *P. syringae* PL3-3 and AT16-2 formed the closest inhibitory/non-inhibitory pair.

Genome mining with antiSMASH revealed a diverse repertoire of BGCs (123 in total) across all analysed *Pseudomonas* strains, varying from 10 (AT16-2) to 23 (P19) identified BGCs per genome (Additional file 6: Table S4). The predicted BGCs belonged to several biosynthetic classes, including NRPS, NRPS-like, RiPP-like, terpene, hydrogen cyanide and siderophore-associated clusters. The proportion of clusters assigned to known BGC types varied among isolates, ranging from 17.6% in *P. fluorescens* PL2-2 to 56.5% in *P. fluorescens* P19. The remaining clusters could not be assigned to a defined BGC type by antiSMASH.

To further compare cluster repertoires across strains, BGCs were grouped into gene cluster families (GCFs) using BiG-SCAPE. The resulting presence–absence matrix revealed 97 different GCFs, of which 18 (including 44 BGCs) were shared by more than one strain, whereas the remaining 79 were found only in a single isolate, i.e. singletons (Additional file 7: Table S5). The distribution of the shared GCFs across strains (Additional file 3: Figure S10) mirrored the whole-genome phylogenetic relationships (Additional file 3: Figure S9). The *P. fluorescens* strains P19, PL2-2, and P22 grouped together, as did the two *P. syringae* and the two *P. putida* strains, whereas *Pseudomonas* sp. PL1-4 formed a separate branch.

Comparative analysis of shared GCFs across strains showed that most were restricted to individual species groups and were present in both inhibitory and non-inhibitory isolates within those groups, suggesting that at least part of the BGC repertoire is retained within related lineages (Figure 5). Only one cluster (PKS-derived arylpolyene cluster, FAM_00002) was detected in inhibitory strains from all three *Pseudomonas* groups (P19, PL2-2, P23 and PL3-3, but not P22), as well as in non-inhibitory *P. syringae* AT16-2. Within the *P. fluorescens* group, all three inhibitory strains (P19, P22 and PL2-2) shared three GCFs (FAM_00001, RiPP-like redox cofactor; FAM_00016, RiPP-like cluster type; FAM_00004, betalactone-type), while P19 and PL2-2 additionally shared three further GCFs (FAM_00005, hydrogen cyanide; FAM_00003, NAGGN-like; FAM_00014, NRPS). Within the *P. putida* group, strain P23 shared a RiPP-like GCF (FAM_00001) with inhibitory strains from the *P. fluorescens* group, while also sharing two other GCFs with its non-inhibitory counterpart P9: an NRP-metallophore/NRPS-like cluster (FAM_00017) and a terpene-precursor cluster (FAM_00011). The highest number of shared clusters was observed within the *P. syringae* group, where most shared GCFs were present in both the inhibitory strain PL3-3 and the non-inhibitory strain AT16-2. Further, we have observed a large variety of singleton GCFs (Table 2) that represent strain-specific biosynthetic potential. We hypothesize that they may reflect relatively recent gain or loss of BGCs, although this remains speculative. Indeed, genomic neighbourhood analysis identified mobility-related genes in the vicinity of only six of the 123 BGCs (4.9%), including both singleton and shared GCFs (Additional file 6: Table S4, Additional file 7: Table S5). These included four singleton GCFs: two in *P. fluorescens* P19 (FAM_00049 and FAM_00078), one in *P. putida* P23 (FAM_00094), and one in *P. syringae* PL3-3 (FAM_00058), as well as two shared GCFs: FAM_00016 in *P. fluorescens* P22 and FAM_00008 in *P. syringae* PL3-3.

**Figure 5.**
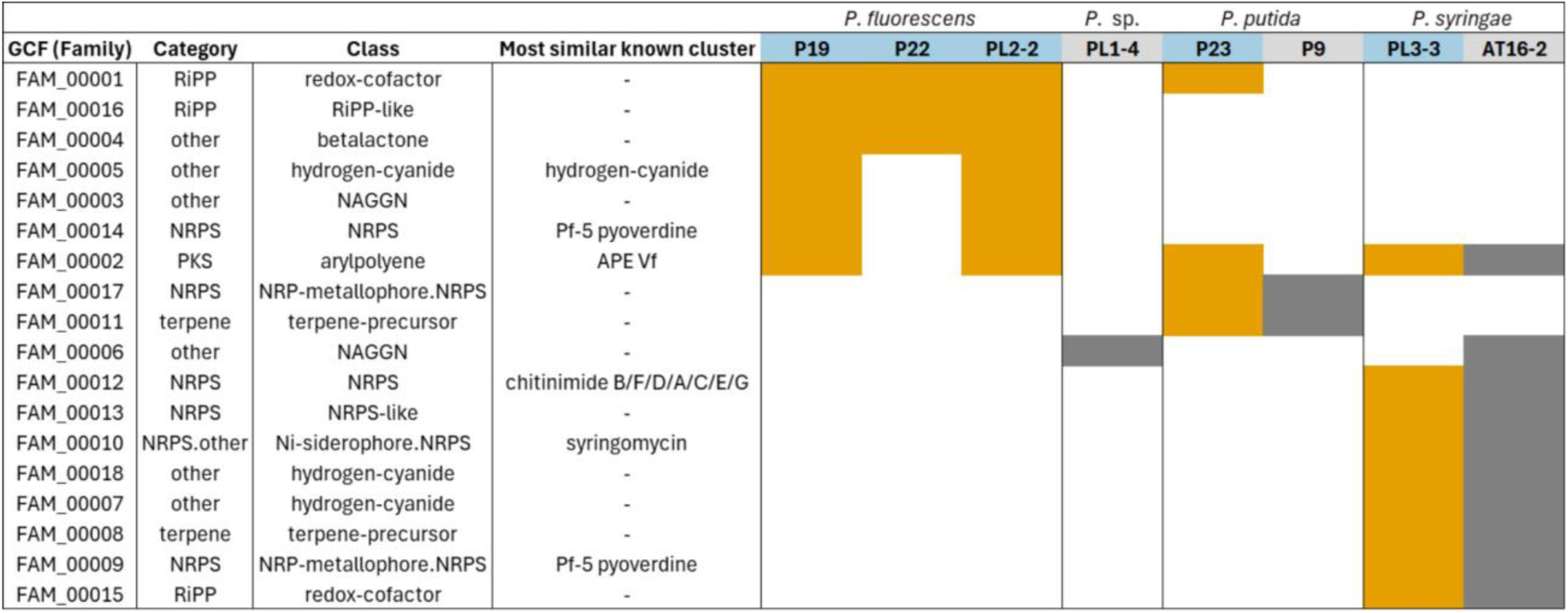
Distribution of shared biosynthetic gene cluster families (GCFs) across *Pseudomonas* isolates. Rows represent individual GCFs grouped by cluster type, and columns represent individual strains. Orange indicates the presence of a given cluster in inhibitory strains, grey indicates presence in non-inhibitory strains, and white indicates absence.

**Table 2.**
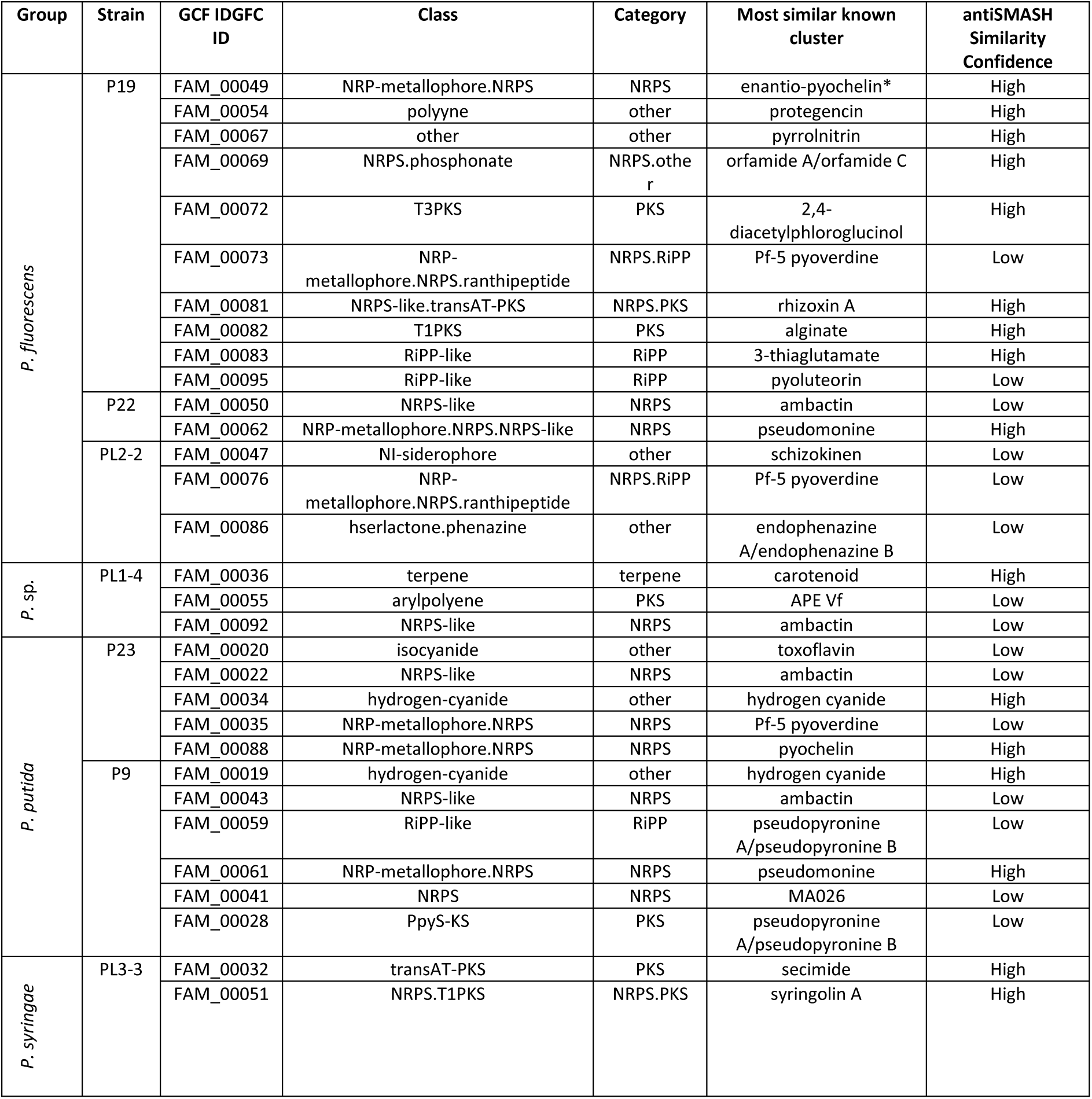
Singleton biosynthetic gene clusters (BGCs) with similarity to previously characterised clusters. Clusters marked with an asterisk indicate the presence of associated mobile genetic elements.

| Group | Strain | GCF IDGFC ID | Class | Category | Most similar known cluster | antiSMASH Similarity Confidence |
|---|---|---|---|---|---|---|
| *P. fluorescens* | P19 | FAM_00049 | NRP-metallophore.NRPS | NRPS | enantio-pyochelin* | High |
|  |  | FAM_00054 | polyyne | other | protegencin | High |
|  |  | FAM_00067 | other | other | pyrrolnitrin | High |
|  |  | FAM_00069 | NRPS.phosphonate | NRPS.other | orfamide A/orfamide C | High |
|  |  | FAM_00072 | T3PKS | PKS | 2,4-diacetylphloroglucinol | High |
|  |  | FAM_00073 | NRP-metallophore.NRPS.ranthipeptide | NRPS.RiPP | Pf-5 pyoverdine | Low |
|  |  | FAM_00081 | NRPS-like.transAT-PKS | NRPS.PKS | rhizoxin A | High |
|  |  | FAM_00082 | T1PKS | PKS | alginate | High |
|  |  | FAM_00083 | RiPP-like | RiPP | 3-thiaglutamate | High |
|  |  | FAM_00095 | RiPP-like | RiPP | pyoluteorin | Low |
|  | P22 | FAM_00050 | NRPS-like | NRPS | ambactin | Low |
|  |  | FAM_00062 | NRP-metallophore.NRPS.NRPS-like | NRPS | pseudomonine | High |
|  | PL2-2 | FAM_00047 | NI-siderophore | other | schizokinen | Low |
|  |  | FAM_00076 | NRP-metallophore.NRPS.ranthipeptide | NRPS.RiPP | Pf-5 pyoverdine | Low |
|  |  | FAM_00086 | hserlactone.phenazine | other | endophenazine A/endophenazine B | Low |
| *P. sp.* | PL1-4 | FAM_00036 | terpene | terpene | carotenoid | High |
|  |  | FAM_00055 | arylpolyene | PKS | APE Vf | Low |
|  |  | FAM_00092 | NRPS-like | NRPS | ambactin | Low |
| *P. putida* | P23 | FAM_00020 | isocyanide | other | toxoflavin | Low |
|  |  | FAM_00022 | NRPS-like | NRPS | ambactin | Low |
|  |  | FAM_00034 | hydrogen-cyanide | other | hydrogen cyanide | High |
|  |  | FAM_00035 | NRP-metallophore.NRPS | NRPS | Pf-5 pyoverdine | Low |
|  |  | FAM_00088 | NRP-metallophore.NRPS | NRPS | pyochelin | High |
|  | P9 | FAM_00019 | hydrogen-cyanide | other | hydrogen cyanide | High |
|  |  | FAM_00043 | NRPS-like | NRPS | ambactin | Low |
|  |  | FAM_00059 | RiPP-like | RiPP | pseudopyronine A/pseudopyronine B | Low |
|  |  | FAM_00061 | NRP-metallophore.NRPS | NRPS | pseudomonine | High |
|  |  | FAM_00041 | NRPS | NRPS | MA026 | Low |
|  |  | FAM_00028 | PpyS-KS | PKS | pseudopyronine A/pseudopyronine B | Low |
| *P. syringae* | PL3-3 | FAM_00032 | transAT-PKS | PKS | secimide | High |
|  |  | FAM_00051 | NRPS.T1PKS | NRPS.PKS | syringolin A | High |

## 4. Discussion

In this study, we investigated antagonism of host-associated pseudomonads towards aquatic oomycetes *A. astaci* and *S. parasitica*, combining *in vitro* phenotypic assays with comparative genomics. Our main findings are: (i) anti-oomycete activity varied substantially among related *Pseudomonas* isolates; (ii) inhibition was consistently stronger against *A. astaci* than against *S. parasitica*; (iii) in most isolates, total inhibition of *A. astaci* was considerably higher than volatile-only inhibition, suggesting a major contribution of non-volatile, presumably diffusible extracellular compoundss, with *P. putida* P23 as an exception with dominantly volatile-mediated inhibition; (iv) siderophore production was detected across isolates but showed no clear relationship with inhibitory activity, suggesting a limited contribution under the tested conditions; and (v) comparative genomic analysis revealed significant variation in BGC repertoires among isolates, with inhibitory strains harbouring candidate clusters with similarity to BGCs of known anti-oomycete compounds. Overall, the obtained results provide a basis for future studies aimed at establishing mechanistic links between inhibition phenotypes and candidate BGCs.

### 4.1. Interspecific differences in susceptibility of aquatic oomycetes to *Pseudomonas* isolates

Antagonistic activity of *Pseudomonas* strains was consistently stronger against *A. astaci* than against *S. parasitica*. A similar pattern of generally higher resistance of *S. parasitica* to external antimicrobial agents has also been reported for plant-derived bioactive compounds [52]. These differences may be at least partly explained by the faster growth rate of *S. parasitica*, which may reduce the effective exposure time of bioactive metabolites on the mycelium during *in vitro* assays, as suggested previously for other oomycetes [53]. However, differences in growth rate likely act together with differences in susceptibility, such as cell surface properties or detoxification capacity. At the same time, the magnitude of inhibition also depended on the bacterial strain, as some *Pseudomonas* isolates inhibited both oomycetes (e.g. *P. fluorescens* PL2-2) whereas others showed activity mainly against *A. astaci* (e.g. *P. putida* P23). Thus, anti-oomycete activity cannot be generalized across pathogens and must instead be empirically verified for each strain–pathogen combination.

### 4.2. High variation of anti-oomycete activity of host-associated *Pseudomonas* isolates

Our results confirm previously reported strong anti-oomycete activity of host-associated *Pseudomonas* isolates [18, 20, 27, 28] but also demonstrate how this activity varies markedly both among and within *Pseudomonas* species groups. The highest number of strongly inhibitory isolates was found within the *P. fluorescens* group in line with previous reports [18–20, 22], whereas isolates affiliated with the *P. putida* and *P. syringae* groups generally showed weaker or less frequent inhibition. This interpretation was supported by the comparative genomic analysis, in which clustering based on BGC composition was broadly consistent with the whole-genome phylogeny, suggesting that antagonistic potential is influenced by broader phylogenetic background. At the same time, isolates belonging to the same *Pseudomonas* species group often exhibited substantial differences in their inhibitory potential, indicating pronounced within-group variation. A similar pattern was previously reported by Orlić et al. [20], where isolates belonging to the same *Pseudomonas* species also showed marked variation in *A. astaci* inhibition. In our dataset, this phenotypic variation coincided with the occurrence of multiple singleton GCFs found in individual inhibitory isolates and absent from their non-inhibitory counterparts, which may reflect relatively recent BGC gain or loss, although this remains hypothetical. If so, this would raise the possibility that at least some inhibitory traits are context dependent and may not be stably maintained in the absence of selection, particularly when associated with metabolically costly pathways or genetically less stable genomic regions [54, 55]. Moreover, antagonistic traits in *Pseudomonas* have been shown to decline or change depending on competitive context, i.e., pathogen presence-absence [56]. However, genomic neighbourhood analysis identified mobility-related genes near only 5% of the detected BGCs, including both singleton and shared GCFs, indicating that mobile-element-associated gain or rearrangement may contribute to BGC diversification in some cases but does not provide a general explanation for the observed strain-specific patterns. Thus, the observed variation in inhibitory activity may also reflect differences in BGC expression or regulation, rather than BGC presence or absence alone.

### 4.3. Potential contribution of diffusible extracellular compounds to inhibition of *A. astaci* by *Pseudomonas* isolates

In most isolates, total inhibition of *A. astaci* was considerably higher than volatile-only inhibition, suggesting that non-volatile, presumably diffusible, compounds produced by *Pseudomonas* strains contributed substantially to inhibition. Genome analyses further revealed a broad repertoire of BGCs potentially encoding diffusible secondary metabolites. These included both clusters with similarity to previously characterised BGCs and uncharacterised clusters, some of which were strain-specific whereas others were shared among multiple inhibitory isolates.

Among the analysed isolates, *P. fluorescens* P19 was particularly notable because it combined the strongest inhibitory phenotype with the highest number of BGCs with similarity to BGCs associated with the production of known diffusible bioactive compounds. Most of these clusters were unique to this strain (singletons), including clusters with similarity to BGCs for the production of pyoluteorin (RiPP-like), rhizoxin (NRPS/PKS-like), pyrrolnitrin, 2,4-diacetylphloroglucinol (DAPG) (T3PKS-like), orfamide (NRPS/phosphonate-like) and 3-thiaglutamate (RiPP-like). All these compounds were previously associated with antimicrobial, antifungal and/or anti-oomycete activity [57–62]. Among these, pyoluteorin is a well-characterised hybrid NRPS/PKS metabolite with demonstrated activity against oomycetes, bacteria and fungi [57, 58]. Each of the other inhibitory isolates also carried its own strain-specific candidate BGCs, such as an endophenazine-like cluster in *P. fluorescens* PL2-2 and a secimide-like cluster in *P. syringae* PL3-3, an isolate that otherwise shared the majority of its BGC repertoire with the non-inhibitory strain AT16-2. Phenazine-producing *Pseudomonas* strains have been shown to inhibit plant-pathogenic oomycetes such as *Phytophthora infestans* and *Pythium* spp. [63–65]. Secimides are glutarimide-containing natural products recently described from *P. syringae* and reported to display selective antifungal activity [66], although anti-oomycete activity has not been demonstrated.

In addition to singleton clusters with similarity to known reference BGCs, inhibitory isolates also shared several uncharacterised clusters that may encode previously undescribed diffusible anti-oomycete metabolites. In particular, the inhibitory *P. fluorescens* isolate P22 contained the highest proportion of uncharacterised BGCs among the sequenced isolates (14 of 17; Additional file 6: Table S4), suggesting that additional, currently unidentified metabolites may contribute to its inhibitory activity. For instance, RiPP-like cluster FAM_00016 was restricted to inhibitory isolates of the *P. fluorescens* group, whereas FAM_00001 was shared by inhibitory isolates from both the *P. fluorescens* and *P. putida* groups, highlighting both clusters as candidates for functional analysis. RiPPs (ribosomally synthesised and post-translationally modified peptides) represent a diverse class of small, typically water-soluble bioactive molecules [67] and are therefore worth considering in future functional studies.

### 4.4. Contribution of volatile compounds to inhibition of *A. astaci* by *Pseudomonas* isolates

Although in most isolates total inhibition was considerably higher than volatile-only inhibition, suggesting a major contribution of non-volatile, presumably diffusible compounds, several *Pseudomonas* isolates exhibited dominant (*P. putida* P23) or partial (*P. fluorescens* PL2-2 and P19, and *P. syringae* PL3-3) *in vitro* inhibition via volatile compounds. In line with this, HCN production was detected in *P. fluorescens* P19 and PL2-2 and *P. putida* P23 on KB medium, while genome analysis identified HCN clusters in all eight isolates. These clusters were diverse and did not group into a single GCF: the HCN-associated BGC in *P. fluorescens* P19 and PL2-2 belonged to the shared family FAM_00005, whereas the remaining HCN-related clusters were assigned to other GCFs. HCN has previously been implicated in anti-oomycete activity, particularly in volatile-mediated inhibition of the plant-pathogenic oomycete *P. infestans in vitro* [68]. However, volatile-mediated inhibition was dominant only in *P. putida* P23, despite the broad occurrence of HCN-related BGCs across strains, suggesting that differences in regulation or metabolite production, rather than BGC presence alone, shape volatile-mediated inhibition. This is supported by the differences in HCN production observed between KB and PG1 media, which indicate that HCN expression is strongly dependent on cultivation conditions, consistent with previous reports [69]. Moreover, because HCN production was assessed only in bacterial monocultures, its actual contribution during bacterial–oomycete interactions under the inhibition-assay conditions cannot be determined directly.

Conversely, the substantial volatile-mediated inhibition exhibited by *P. syringae* PL3-3 despite only weak HCN production indicates that other volatile metabolites may also contribute to antagonistic activity. Consistent with this interpretation, several gene clusters related to lactones and terpene precursors were identified and may be associated with the production of volatile organic compounds and anti-oomycete activity [70, 71]. More broadly, HCN represents only one component of the complex *Pseudomonas* volatile metabolome, and multiple VOCs may contribute jointly to antagonistic interactions [68, 72].

Finally, although the antagonistic relevance of HCN and other volatiles in aquatic systems cannot be demonstrated by plate assays performed in this study, they can diffuse in both gas and water phases and may therefore contribute to short-range chemical interactions in aquatic environments under suitable microenvironmental conditions [72]. Their relative contribution in this model system therefore requires further experimental testing (see Section 4.6).

### 4.5. Role of iron availability and siderophores in inhibition of *A. astaci* by *Pseudomonas* isolates

In contrast to previous reports suggesting that siderophores may contribute to oomycete inhibition via iron competition or direct toxicity [6, 23, 73], our results did not support a clear association between siderophore production and inhibitory phenotype. Siderophore activity was detected in isolates across species groups, irrespective of their antagonistic activity toward *A. astaci*. Also, it depended on environmental conditions, i.e. it was more pronounced under iron-limited conditions, consistent with their established role in iron acquisition in *Pseudomonas* [57, 66], and in KB compared to PG1 medium. Further, genome analysis showed that all isolates possessed one or more siderophore BGCs, including clusters with similarity to known BGCs linked to pyoverdine, pseudomonine, enantio-pyochelin and schizokinen production. Among these, pyoverdine is the siderophore most often implicated in oomycete inhibition through iron competition [6, 73, 74]. Putative pyoverdine-related BGCs were identified in several isolates, including two clusters in each of the inhibitory *P. fluorescens* isolates P19 and PL2-2, both of which also exhibited pyoverdine-associated fluorescence. Overall, our results do not support siderophore production as a general determinant of *A. astaci* inhibition, although pyoverdine or other siderophores may contribute to antagonism in a strain-and condition-dependent manner.

### 4.6. Study limitations and future directions

This study has several limitations that also point to directions for future research. First, the comparative genomic analyses describe only the biosynthetic potential of the strains. A BGC similar to a characterised cluster does not by itself show which compound it encodes (similar clusters may produce related or different metabolites), nor whether this compound is produced or contributes to inhibition. Moreover, with only eight sequenced isolates, the comparisons between BGC content and inhibitory phenotypes lack statistical power and remain mainly descriptive. The contribution of the candidate BGCs discussed above should therefore be tested in a larger set of sequenced strains, combining metabolomic profiling with targeted BGC deletions.

Second, zoospores are the main dispersal and infective stages of oomycetes, and germinating cysts penetrate the host. As most of our isolates originate from fish or crayfish, where they are part of the healthy host biofilm, they are likely to encounter zoospores and germinating cysts at this important site of host defence. However, we analysed only the inhibition of mycelial growth, the stage that causes disease symptoms within host tissues, so it remains to be tested whether *Pseudomonas* isolates or their metabolites also affect zoospore production, motility, encystment or germination.

Third, the split-plate assay quantified total and volatile-only inhibition; the contribution of non-volatile, presumably diffusible, compounds was inferred from their difference and should be confirmed directly, e.g. by testing cell-free culture supernatants or extracts of the isolates against the oomycetes.

Finally, translating these in vitro findings into the protection of crayfish and fish against oomycete diseases remains a substantial step. In natural and aquaculture settings bacteria, oomycetes and their hosts are exposed to fluctuating environmental conditions, such as daily and seasonal changes in water temperature and available nutrients, which affect each of them individually as well as their interactions [75]. The dependence of bacterial phenotypes on nutrient availability was evident already in vitro: in the bacteria-only phenotypic assays, HCN production and siderophore activity of the isolates differed between media. It is therefore impossible to extrapolate the inhibition results, which were obtained in co-culture assays performed on a single medium (PG1) and at a single temperature (18 °C), to natural or aquaculture environments. Accordingly, the terms inhibitory and non-inhibitory, as used throughout this study, refer (only) to the tested assay conditions, and the inhibitory activity of the isolates may differ on other media, under other temperature regimes and in the presence of the host. In particular, it remains to be tested whether the isolates retain their inhibitory activity at lower temperatures (e.g., 10–15 °C), which are common in the natural habitats and aquaculture systems of crayfish and fish. Beyond such in vitro tests, the most promising isolates should ultimately be evaluated in animal infection trials, including host colonisation and safety for the host and environment, before any application in aquaculture.

## 5. Conclusions and future perspectives

In conclusion, our findings do not support a single universal metabolite or conserved biosynthetic pathway that would explain anti-oomycete activity of the investigated *Pseudomonas* isolates. Instead, they are consistent with a scenario in which anti-oomycete activity may arise from different combinations of known and uncharacterised biosynthetic pathways. Among the analysed isolates, *P. fluorescens* P19 showed the closest correspondence between phenotype (strong oomycete inhibition, as well as HCN and siderophore production) and biosynthetic potential: besides HCN-and siderophore-related BGCs, it contained clusters exclusive to inhibitory strains (e.g. RiPP-like clusters) and those with similarity to characterised bioactive pathways (e.g. pyoluteorin, rhizoxin, pyrrolnitrin, DAPG and orfamide). These BGCs represent promising candidates for future functional validation. Together, these results narrow down the candidate biosynthetic pathways potentially associated with anti-oomycete activity and highlight the value of combining phenotypic assays with comparative genomics to identify both known and previously uncharacterised inhibitory metabolites.

## Abbreviations

BGC: Biosynthetic gene cluster
CAS: Chrome Azurol S
DAPG: 2,4-Diacetylphloroglucinol
GCF: Gene cluster family
HCN: Hydrogen cyanide
KB: King’s B medium
NRPS: Non-ribosomal peptide synthetase
OD600: Optical density at 600 nm
PKS: Polyketide synthase
RiPP: Ribosomally synthesised and post-translationally modified peptide

## Ethics approval and consent to participate

Not applicable.

## Consent for publication

Not applicable.

## Availability of data and materials

The genome assemblies generated in this study have been deposited in the NCBI BioProject database under accession number PRJNA1494142 (https://www.ncbi.nlm.nih.gov/bioproject/PRJNA1494142); the accession numbers of the individual genome assemblies are provided in Table 1. The 16S rRNA gene sequences are available in GenBank under accession numbers PP535706–PP536037 (Additional file 2: Table S1). The datasets supporting the remaining conclusions of this article are included within the article and its additional files.

## Authors’ contributions

**Ela Šarić:** Formal analysis, Data curation, Writing – original draft, Writing – review & editing, Visualization.

**Anđela Miljanović:** Investigation, Formal analysis, Data curation.

**Petra Struški:** Methodology, Investigation, Formal analysis, Data curation, Visualization.

**Simone Oberhaensli**: Methodology, Investigation, Formal analysis, Data curation, Writing – review & editing.

**Livia Jerjen:** Methodology, Investigation, Formal analysis, Writing – review & editing.

**Jurica Žučko:** Methodology, Formal analysis, Writing – review & editing, Visualization.

**Heike Schmidt-Posthaus:** Writing – review & editing, Resources.

**Dora Pavić:** Formal analysis, Data curation.

**Ivana Maguire:** Investigation, Writing – review & editing.

**Julius Hermanns:** Investigation, Writing – review & editing.

**Laure Weisskopf:** Methodology, Writing – review & editing, Resources.

**Tobia Pretto:** Investigation, Resources.

**Ana Bielen:** Conceptualization, Methodology, Writing – original draft, Writing – review & editing, Resources, Supervision, Project administration, Funding acquisition.

All authors have read and approved the final version of the manuscript and agree to its submission.

## Acknowledgements

The authors are thankful to Maja Valentić for the technical assistance during laboratory work. We also thank Leo Barišić, Matilda Majerščak-Škorlić, Elena Pirović and Lucija Vukšić for their assistance in parts of the experimental work. We thank the Next Generation Sequencing Platform of the University of Bern for performing the high-throughput sequencing experiments. Calculations were performed on UBELIX (https://www.id.unibe.ch/hpc), the HPC cluster at the University of Bern.

## AI declaration

During the preparation of this manuscript, the authors used Claude Opus 5.5 (Anthropic) for language editing and for assistance with renumbering of references. Figure 1 was prepared in Microsoft PowerPoint with the assistance of Claude, based on the study design and content provided by the authors. The authors reviewed and verified all AI-assisted content and take full responsibility for the content of this publication.

## Additional files

**Additional file 1 (DOCX).** Sampling of host-associated bacteria and taxonomic composition of the isolate collection. Describes the sampling and culturing of the host-associated bacterial isolates and the preliminary identification of the isolates, and contains the sampling summary (Table A1), the proportions of the most prevalent bacterial genera isolated from crayfish and fish (Figure A1), and the taxonomic composition of the bacterial communities associated with crayfish (Table A2) and with fish (Table A3).

**Additional file 2 (XLSX).** Table S1. List of bacterial isolates used in this study, including isolate codes, host species, organ of origin, taxonomic identification based on 16S rRNA gene sequencing, percentage identity, and GenBank accession numbers.

**Additional file 3 (DOCX).** Figures S1–S10. Figure S1, representative plate co-culture inhibition assay; Figure S2, representative split-plate assay; Figure S3, total and volatile-only inhibition of oomycete growth by the eight selected isolates; Figure S4, qualitative assessment of hydrogen cyanide (HCN) production on KB and PG1 media; Figure S5, qualitative assessment of siderophore production using the CAS assay on solid media; Figure S6, quantification of siderophore activity using the liquid CAS assay under iron-limited (Fe−) and iron-supplemented (Fe+) conditions; Figure S7, qualitative assessment of broad-range fluorescence under UV illumination; Figure S8, quantification of pyoverdine-associated fluorescence in liquid KB and PG1 media; Figure S9, whole-genome phylogenetic placement of the eight selected isolates relative to reference genomes; Figure S10, clustering of the selected isolates based on shared biosynthetic gene cluster family (GCF) composition.

**Additional file 4 (XLSX).** Table S2. List of all *Pseudomonas* isolates used in this study, including inhibition zones against *Aphanomyces astaci* and *Saprolegnia parasitica*.

**Additional file 5 (XLSX).** Table S3. Raw measurements and summary statistics (mean ± SD) for co-culture inhibition, split-plate assays, siderophore production, and pyoverdine-associated fluorescence used for the statistical analyses presented in the main text.

**Additional file 6 (XLSX).** Table S4. List of biosynthetic gene clusters (BGCs) predicted by antiSMASH in the analysed *Pseudomonas* isolates, including associated mobile genetic elements.

**Additional file 7 (XLSX).** Table S5. Distribution of biosynthetic gene cluster families (GCFs) identified by BiG-SCAPE across the analysed *Pseudomonas* isolates, including BGC category and class, genomic region, and singleton status.

