## Supplementary material for "Variation of anti-oomycete activity in *Pseudomonas* spp.: phenotypic characterization and comparative genomics": Additional file 1.docx

**Additional file 1. Sampling of host-associated bacteria and taxonomic composition of the isolate collection**

This additional file describes the sampling of apparently healthy trout and crayfish, the culturing of the host-associated bacterial isolates, their identification by 16S rRNA gene sequencing, and the taxonomic composition of the resulting isolate collection. In the main text, this collection was used solely as a source of *Pseudomonas* isolates; isolates were not compared according to sample origin, host species or sampling date.

**1. Sampling and culturing of host-associated bacterial isolates**

We have collected skin and exoskeleton biofilm samples of apparently healthy trout and crayfish, respectively, to culture host-associated bacterial isolates. Crayfish originated from natural rivers and lakes and were trapped using baited Li–Ni traps [1], while trout individuals were obtained from a commercial fish farm (Table A1).

The crayfish cuticle-associated community was sampled according to Pavić et al. [2]. Fish skin was sampled using sterile swabs, and the collected material was suspended in 1 mL of sterile PBS (140 mM NaCl, 2.68 mM KCl, 3.81 mM Na_2_HPO_4_ × 2 H_2_O, 1.47 mM KH_2_PO_4_). Suspensions were serially diluted from 10^-1^ to 10^-3^, plated on LB agar, and incubated at 21 °C. Distinct colony morphotypes observed after 48 and 72 h were purified by successive streaking and preserved in 30% glycerol at − 80 °C. A total of 168 and 135 bacterial isolates from the surface of fish and crayfish were collected, respectively. This collection was supplemented with additional 25 *Pseudomonas* isolates originating from different fish and crayfish organs collected during the routine diagnostic work at the Istituto Zooprofilattico Sperimentale delle Venezie, Legnaro, Italy. All isolates are listed in Additional file 2: Table S1.

**2. Preliminary identification of bacterial isolates by 16S rRNA gene sequencing**

To grow the biomass for genomic DNA isolation, 100 μL of glycerol suspension of each isolate was inoculated onto LB solid medium and incubated at 21 °C overnight. Next, the biomass was scraped of the plate surface and total genomic DNA was isolated using NucleoSpin Microbial DNA kit (Macherey-Nagel) following the protocol for Gram-negative bacteria. Cells were lysed with Macherey-Nagel type B beads by vortexing on a Corning® LSE™ device for 12 minutes at medium power. DNA concentrations were between 20 and 300 ng μL^-1^, as measured with a Quantus fluorometer (Promega) and QuantiFluor ONE dsDNA System dye (Promega).

Nearly full-length 16S rRNA gene fragments were amplified using primers 27F and 1492R [3], following the PCR mixture and cycling conditions described by Orlić et al. [4]. Amplicons were purified and sequenced bidirectionally by Microsynth, Austria. Chromatograms were assembled and edited in GeneStudio, and genus-level assignments were obtained by BLASTN comparison against the NCBI 16S rRNA database.

**Table A1.** Sampling summary.

| **Host species** | **Number of sampled animals** | **Location** | **Coordinates** | **Sampling date** |
| --- | --- | --- | --- | --- |
| Signal crayfish (*Pacifastacus leniusculus*) | 10 | Korana River (Ladvenjak) | N 45°24'44"  E 15°36'30" | 22.10.2021. |
| Narrow-clawed crayfish (*Pontastacus leptodactylus*) | 10 | Korana River (Ušće) | N 45°27'54"  E 15°33'58" | 22.10.2021. |
| Noble crayfish (*Astacus astacus*) | 10 | Plitvice lakes (Novakovića Brod) | N 44°54'7"  E 15°36'39" | 17.09.2021. |
| White-clawed crayfish (*Austropotamobius fulcisianus*) | 6 | Jugovski potok | N 45°25'20"  E 13°47'3" | 17.09.2021. |
| Stone crayfish (*Austropotamobius torrentium*) | 10 | Prijeboj | N 44°49'50"  E 15°39'49" | 17.09.2021. |
| Brown trout (*Salmo trutta*) | 10 | fish farm Gračani V.I.R.I.B.U.S. | N 45° 51' 24.20858"  E 15° 57' 52.65883" | 25.11.2021. |
| Rainbow trout (*Oncorhynchus mykiss*) | 11 |  |  |  |

**3. Taxonomic composition of the isolate collection**

In total, 135 isolates were collected from five crayfish species and 168 from two trout species (Additional file 2: Table S1). Most isolates from the crayfish cuticle belonged to the phylum Pseudomonadota (65.2%), followed by Bacteroidota (18.5%), Bacillota (14.1%) and Actinobacteriota (2.2%) (Table A2). The most prevalent genera were *Pseudomonas* (27.4%), *Aeromonas* (19.3%), *Flavobacterium* (11.9%), *Chryseobacterium* (6.7%) and *Exiguobacterium* (6.7%) (Figure A1, Table A2). Among the bacterial isolates collected from the skin of trout, Pseudomonadota (45.2%) and Actinobacteriota (36.3%) were the dominant phyla, followed by Bacteroidota (16.1%), Bacillota (1.8%) and Deinococcota (0.6%) (Table A3). The most common genera were *Paracoccus* (17.3%), *Microbacterium* (12.5%), *Massilia* (8.3%), *Arthrobacter* (8.3%) and *Psychrobacter* (7.7%) (Figure A1).

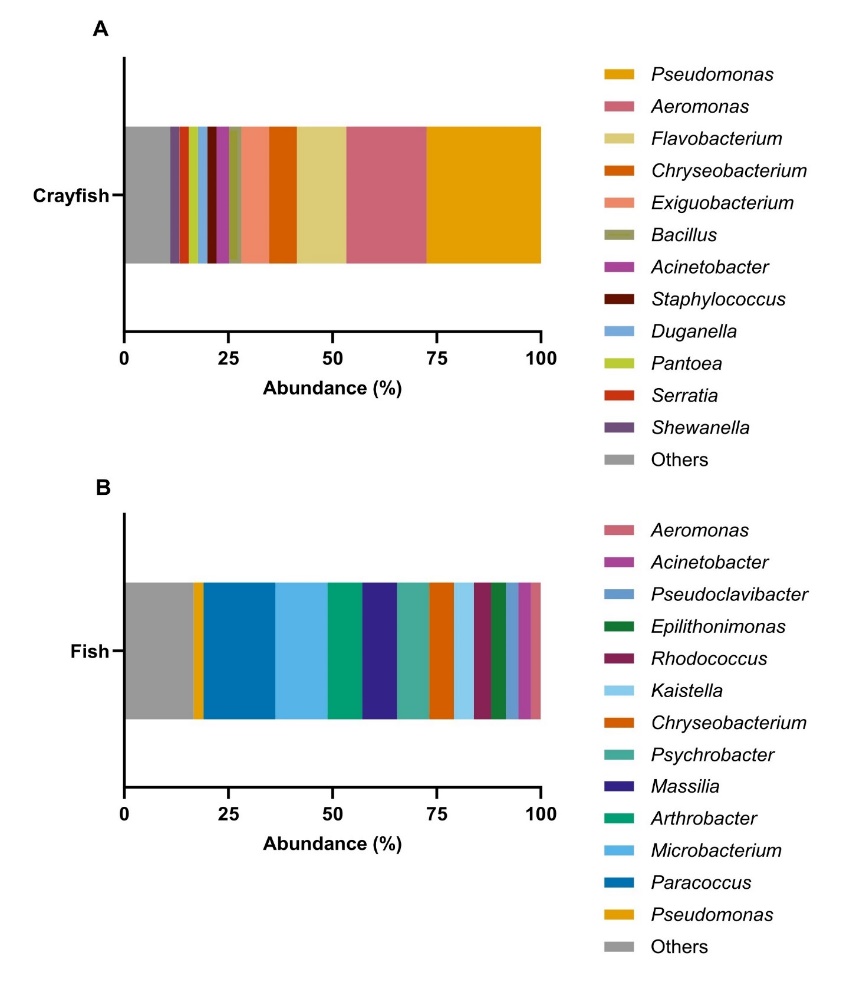
<https://doi.org/10.1128/jb.173.2.697-703.1991>

**Figure A1.** Proportion of isolates (%) belonging to the most prevalent bacterial genera isolated from crayfish (A) and fish (B). Genera representing less than 2% of the isolates were grouped under "Others".

**Table A2.** Taxonomic composition of the bacterial communities associated with different crayfish species. AA - *Astacus astacus*, POL - *Pontastacus leptodactylus*, AF - *Austropotamobius fulcisianus*, AT - *Austropotamobius torrentium*, PL *- Pacifastacus leniusculus*. Cell shading is proportional to the number of isolates.

| **Phylum** | **Genus** | **Number of isolates from each crayfish species** | | | | | **Number of isolates for individual genus/total isolates number** |
| --- | --- | --- | --- | --- | --- | --- | --- |
|  |  | **AA** | **POL** | **AF** | **AT** | **PL** |  |
| **Actinobacteriota** | *Curtobacterium* | 0 | 1 | 0 | 0 | 0 | 1/135 |
|  | *Frigoribacterium* | 0 | 0 | 0 | 0 | 1 | 1/135 |
|  | *Microbacterium* | 0 | 0 | 0 | 1 | 0 | 1/135 |
| **Bacillota** | *Bacillus* | 0 | 0 | 3 | 1 | 0 | 4/135 |
|  | *Exiguobacterium* | 0 | 0 | 2 | 1 | 6 | 9/135 |
|  | *Planomicrobium* | 0 | 0 | 0 | 0 | 1 | 1/135 |
|  | *Psychrobacillus* | 0 | 0 | 0 | 0 | 1 | 1/135 |
|  | *Rossellomorea* | 0 | 0 | 1 | 0 | 0 | 1/135 |
|  | *Staphylococcus* | 0 | 0 | 3 | 0 | 0 | 3/135 |
| **Bacteroidota** | *Chryseobacterium* | 0 | 5 | 1 | 2 | 1 | 9/135 |
|  | *Flavobacterium* | 1 | 3 | 0 | 8 | 4 | 16/135 |
| **Pseudomonadota** | *Acinetobacter* | 2 | 1 | 0 | 1 | 0 | 4/135 |
|  | *Aeromonas* | 4 | 7 | 5 | 7 | 3 | 26/135 |
|  | *Duganella* | 0 | 1 | 0 | 0 | 2 | 3/135 |
|  | *Erwinia* | 0 | 1 | 1 | 0 | 0 | 2/135 |
|  | *Ewingella* | 0 | 1 | 0 | 0 | 0 | 1/135 |
|  | *Hafnia* | 0 | 0 | 0 | 1 | 0 | 1/135 |
|  | *Obesumbacterium* | 0 | 0 | 1 | 1 | 0 | 2/135 |
|  | *Pantoea* | 0 | 0 | 1 | 0 | 2 | 3/135 |
|  | *Pseudomonas* | 1 | 13 | 5 | 9 | 9 | 37/135 |
|  | *Rahnella* | 1 | 0 | 0 | 0 | 0 | 1/135 |
|  | *Serratia* | 0 | 0 | 0 | 3 | 0 | 3/135 |
|  | *Shewanella* | 1 | 0 | 0 | 0 | 2 | 3/135 |
|  | *Stenotrophomonas* | 0 | 1 | 0 | 0 | 0 | 1/135 |
|  | *Yersinia* | 1 | 0 | 0 | 0 | 0 | 1/135 |

**Table A3.** Taxonomic composition of the bacterial communities associated with different fish species. ST – *Salmo trutta*, OM – *Oncorhynchus mykiss*. Cell shading is proportional to the number of isolates.

| **Phylum** | **Genus** | **Number of isolates from each fish species** | | **Number of isolates for individual genus/total isolates number** |
| --- | --- | --- | --- | --- |
|  |  | **ST** | **OM** |  |
| **Actinobacteriota** | *Agrococcus* | 1 | 0 | 1/168 |
|  | *Arthrobacter* | 9 | 5 | 14/168 |
|  | *Brachybacterium* | 0 | 1 | 1/168 |
|  | *Clavibacter* | 1 | 0 | 1/168 |
|  | *Curtobacterium* | 0 | 2 | 2/168 |
|  | *Dietzia* | 1 | 0 | 1/168 |
|  | *Kocuria* | 0 | 3 | 3/168 |
|  | *Labedella* | 0 | 1 | 1/168 |
|  | *Leucobacter* | 0 | 1 | 1/168 |
|  | *Microbacterium* | 13 | 8 | 21/168 |
|  | *Pseudoclavibacter* | 3 | 2 | 5/168 |
|  | *Rhodococcus* | 5 | 2 | 7/168 |
|  | *Rothia* | 1 | 2 | 3/168 |
| **Bacillota** | *Bacillus* | 0 | 1 | 1/168 |
|  | *Macrococcus* | 0 | 2 | 2/168 |
| **Bacteroidota** | *Chryseobacterium* | 4 | 6 | 10/168 |
|  | *Epilithonimonas* | 1 | 5 | 6/168 |
|  | *Flavobacterium* | 0 | 2 | 2/168 |
|  | *Kaistella* | 4 | 4 | 8/168 |
|  | *Pedobacter* | 1 | 0 | 1/168 |
| **Deinococcota** | *Deinococcus* | 1 | 0 | 1/168 |
| **Pseudomonadota** | *Acinetobacter* | 2 | 3 | 5/168 |
|  | *Aeromonas* | 1 | 3 | 4/168 |
|  | *Buttiauxella* | 0 | 1 | 1/168 |
|  | *Hafnia* | 0 | 1 | 1/168 |
|  | *Kluyvera* | 0 | 1 | 1/168 |
|  | *Massilia* | 11 | 3 | 14/168 |
|  | *Melaminivora* | 1 | 0 | 1/168 |
|  | *Obesumbacterium* | 0 | 1 | 1/168 |
|  | *Paracoccus* | 16 | 13 | 29/168 |
|  | *Pseudomonas* | 4 | 0 | 4/168 |
|  | *Psychrobacter* | 5 | 8 | 13/168 |
|  | *Sphingomonas* | 1 | 1 | 2/168 |
