## Supplementary material for "Variation of anti-oomycete activity in *Pseudomonas* spp.: phenotypic characterization and comparative genomics": Additional file 3.docx

**Additional file 3. Supplementary Figures S1–S10**

| 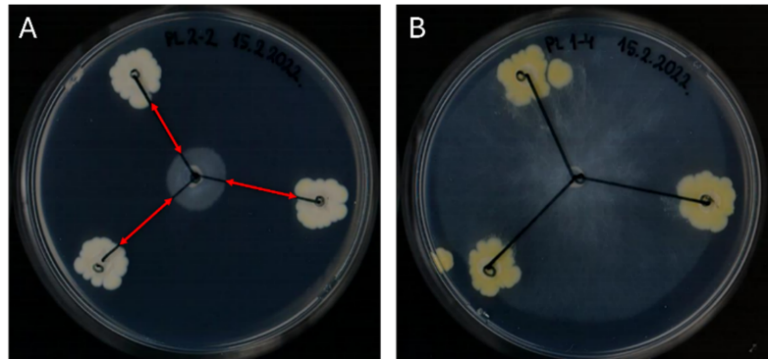 | | **Figure S1.** *A. astaci* mycelial growth in the presence of two *Pseudomonas* strains using plate-based co-culture inhibition assay: (A) inhibiting isolate *P.* *fluorescens* PL2-2 (mycelial growth is almost completely stopped), and (B) *Pseudomonas* sp. PL1-4 (mycelial growth undisturbed). Red arrows are inhibition zones, i.e., distances between the edge of the bacterial colony and the edge of the mycelium. |
| --- | --- | --- |
| 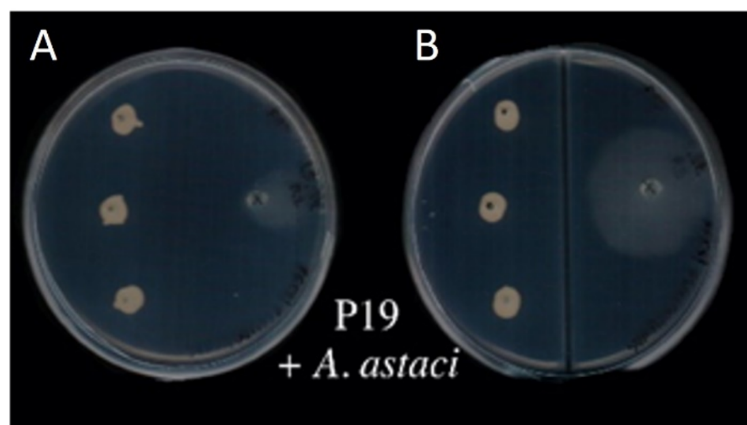 | **Figure S2.** *A. astaci* mycelial growth in the presence of *P. fluorescens* P19 using split-plate assay: (A) inhibition on a regular plate, where mycelial growth is almost completely suppressed, and (B) inhibition on a split plate, where mycelial growth remains largely unaffected. | |

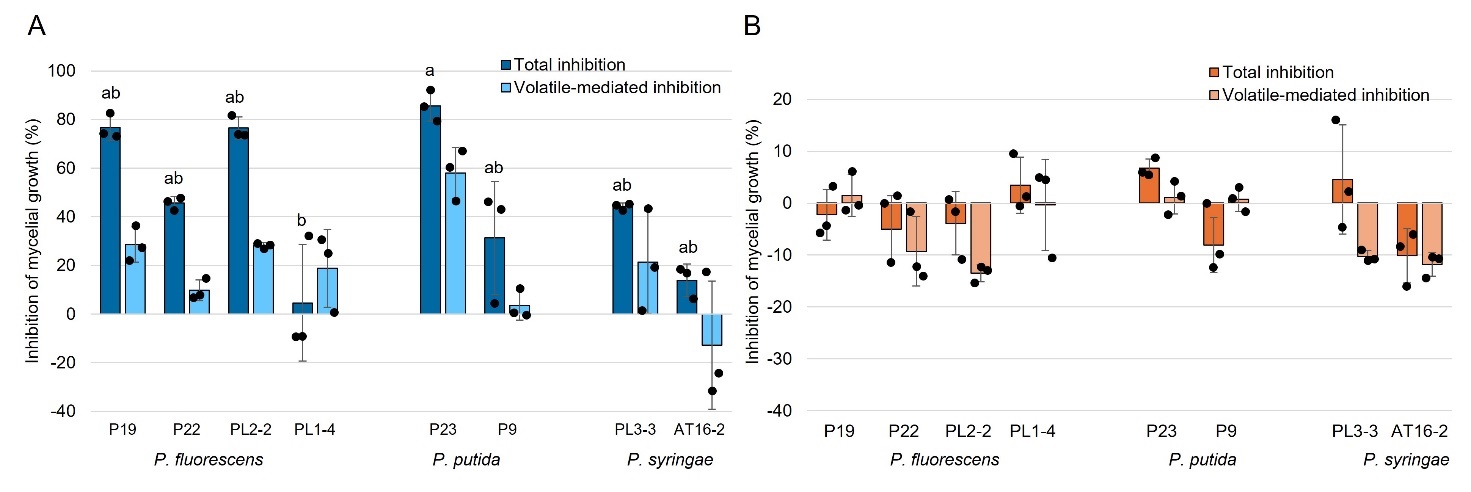
**Figure S3. Total and volatile-only inhibition of (A) *A. astaci* and (B) *S. parasitica* by eight selected *Pseudomonas* isolates.** (A) Dark blue and (B) dark orange bars represent total inhibition, while (A) light blue and (B) light orange bars represent volatile-only inhibition. Values represent mean ± standard deviation (*n* = 3), with individual measurements shown as black dots. Different letters indicate statistically significant differences among isolates in total inhibition of *A. astaci* (Dunn’s post hoc test with Holm correction, *p* < 0.05). Isolates sharing at least one letter are not significantly different. No significant pairwise differences were detected in volatile-only inhibition of *A. astaci* or in either inhibition measure for *S. parasitica* after Holm correction.

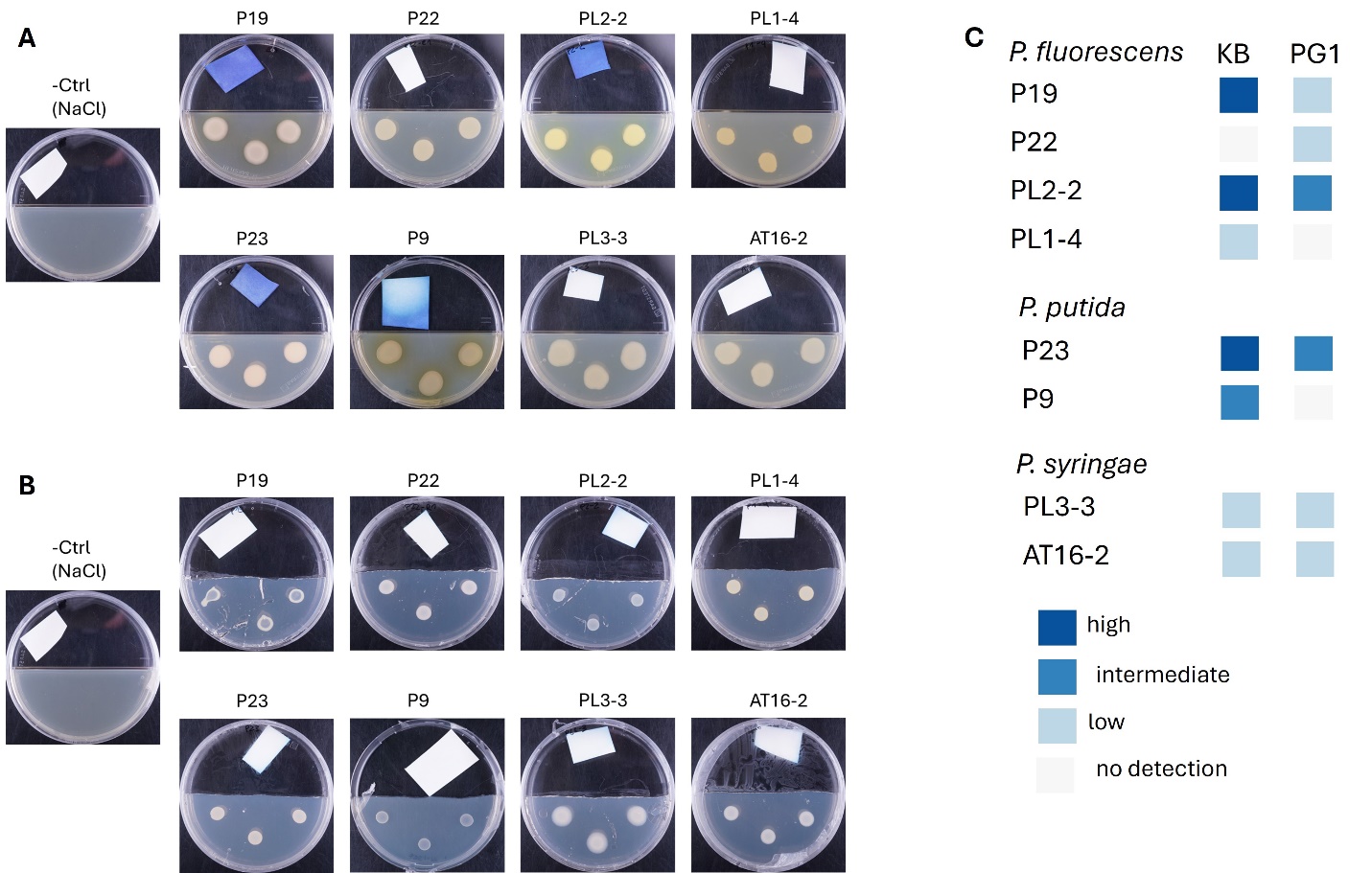

**Figure S4. Qualitative assessment of hydrogen cyanide (HCN) production by the analysed *Pseudomonas* isolates grown on KB and PG1 media.** Representative cyanide detection assays performed on (A) KB and (B) PG1 media are shown. HCN production was assessed using HCN detection paper placed in the agar-free compartment of the Petri dish and scored visually according to the intensity of blue discoloration after 48 h of incubation. Panel C summarizes the qualitative HCN production observed for each isolate on both media, classified into four categories (high, intermediate, low and no detection).

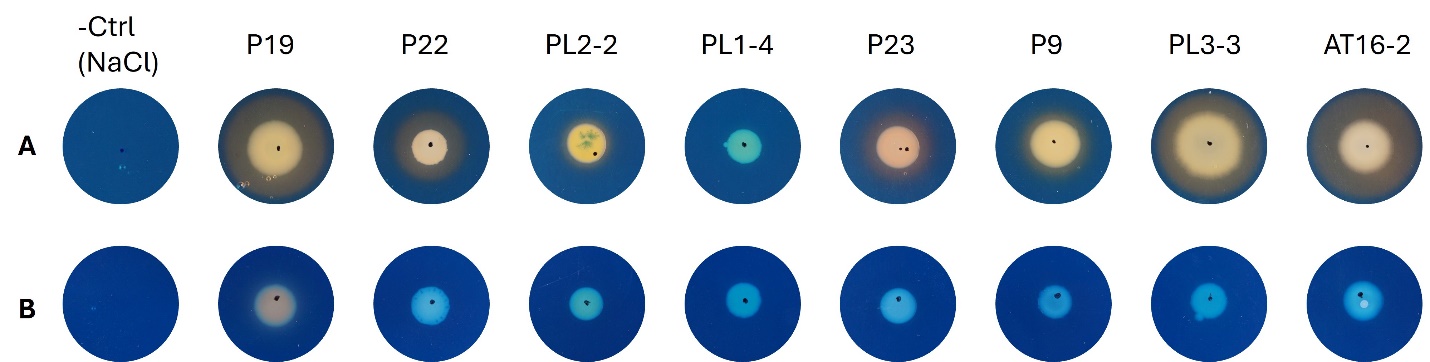

**Figure S5. Qualitative detection of siderophore production by eight selected *Pseudomonas* isolates using CAS assay.** Isolates were grown on (A) KB and (B) PG1 agar media. Orange-yellow halos surrounding bacterial colonies indicate siderophore activity. Uninoculated media (−Ctrl) were used as negative controls.

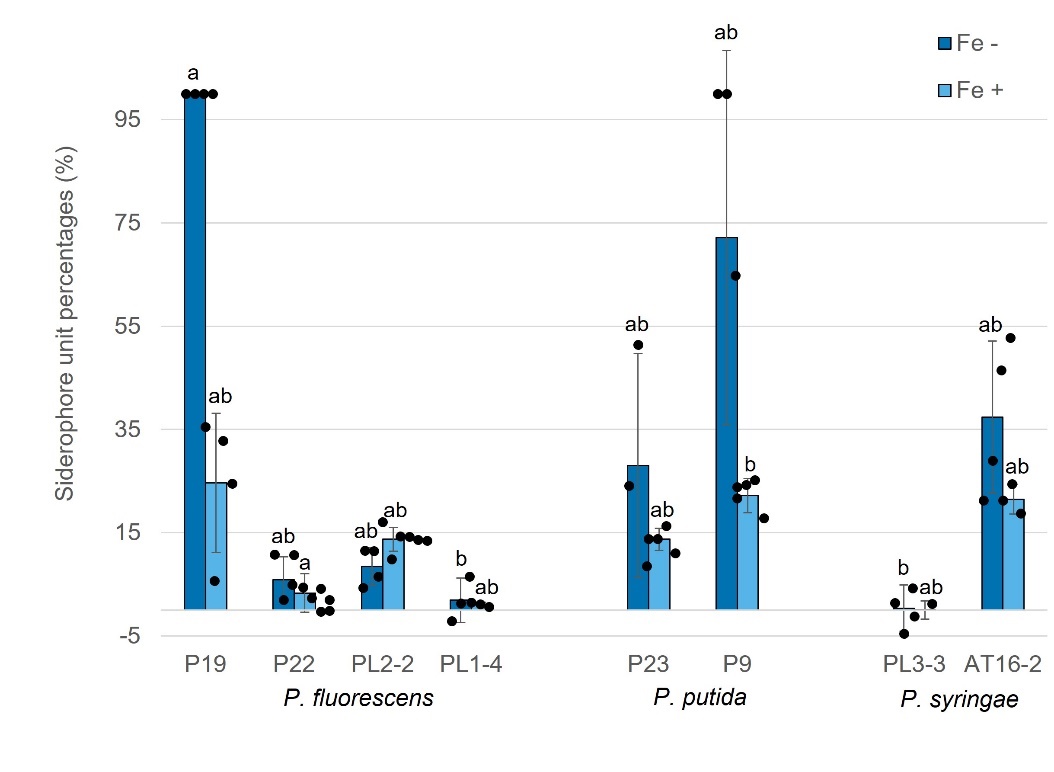

**Figure S6. Quantitative assessment of siderophore production by eight selected *Pseudomonas* isolates using the CAS liquid assay in PG1 medium.** Siderophore activity was measured under iron-limited (Fe−) and iron-supplemented (Fe+) conditions and expressed as percentage siderophore units. Data are presented as mean ± SD (*n* = 2–7), with individual biological replicates shown as black dots. Different letters indicate statistically significant differences among isolates within the same iron condition (Dunn’s post hoc test with Holm correction, *p* < 0.05). Isolates sharing at least one letter are not significantly different.

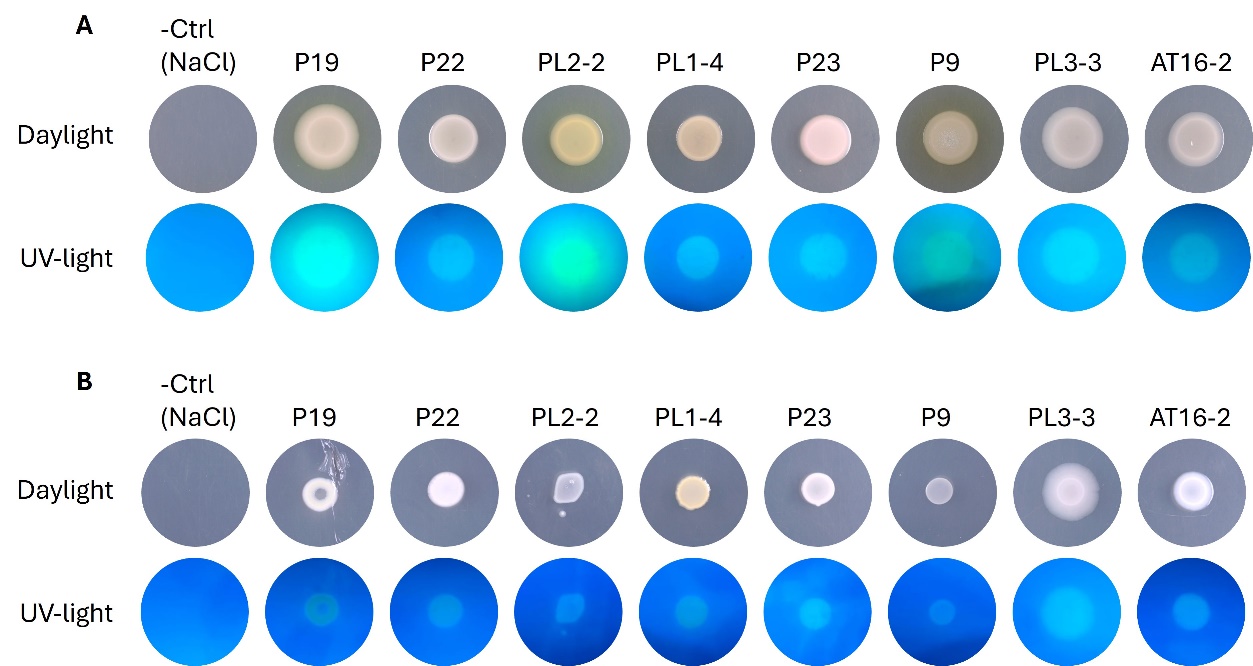

**Figure S7. Qualitative assessment of broad-range fluorescence emitted by the analysed *Pseudomonas* isolates grown on (A) KB and (B) PG1 solid media.** Colonies are shown under visible light and UV illumination. Broad-range fluorescence was assessed qualitatively based on the presence or absence of fluorescence under UV light.

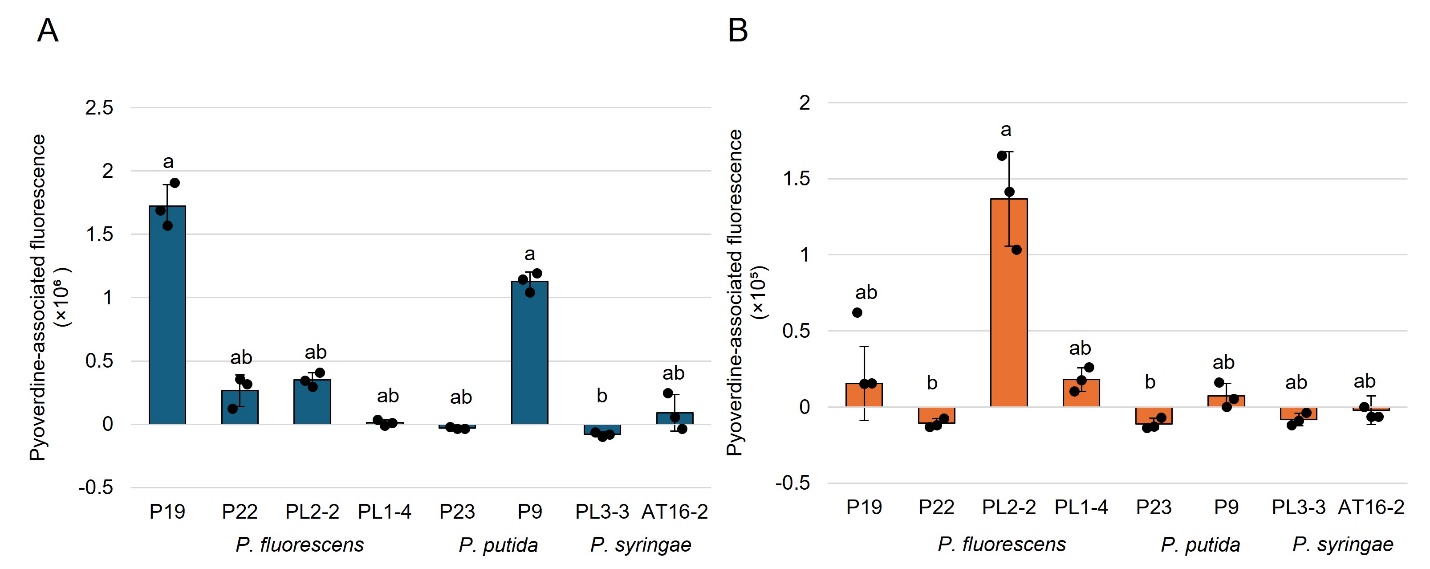

**Figure S8. Quantification of pyoverdine-associated fluorescence of the analysed *Pseudomonas* isolates grown in liquid KB (A) and PG1 (B) media.** Bars represent mean ± SD (*n* = 3), with individual biological replicates shown as black dots. Different letters indicate statistically significant differences among isolates within the same growth medium (Dunn’s post hoc test with Holm correction, *p* < 0.05). Isolates sharing at least one letter are not significantly different.

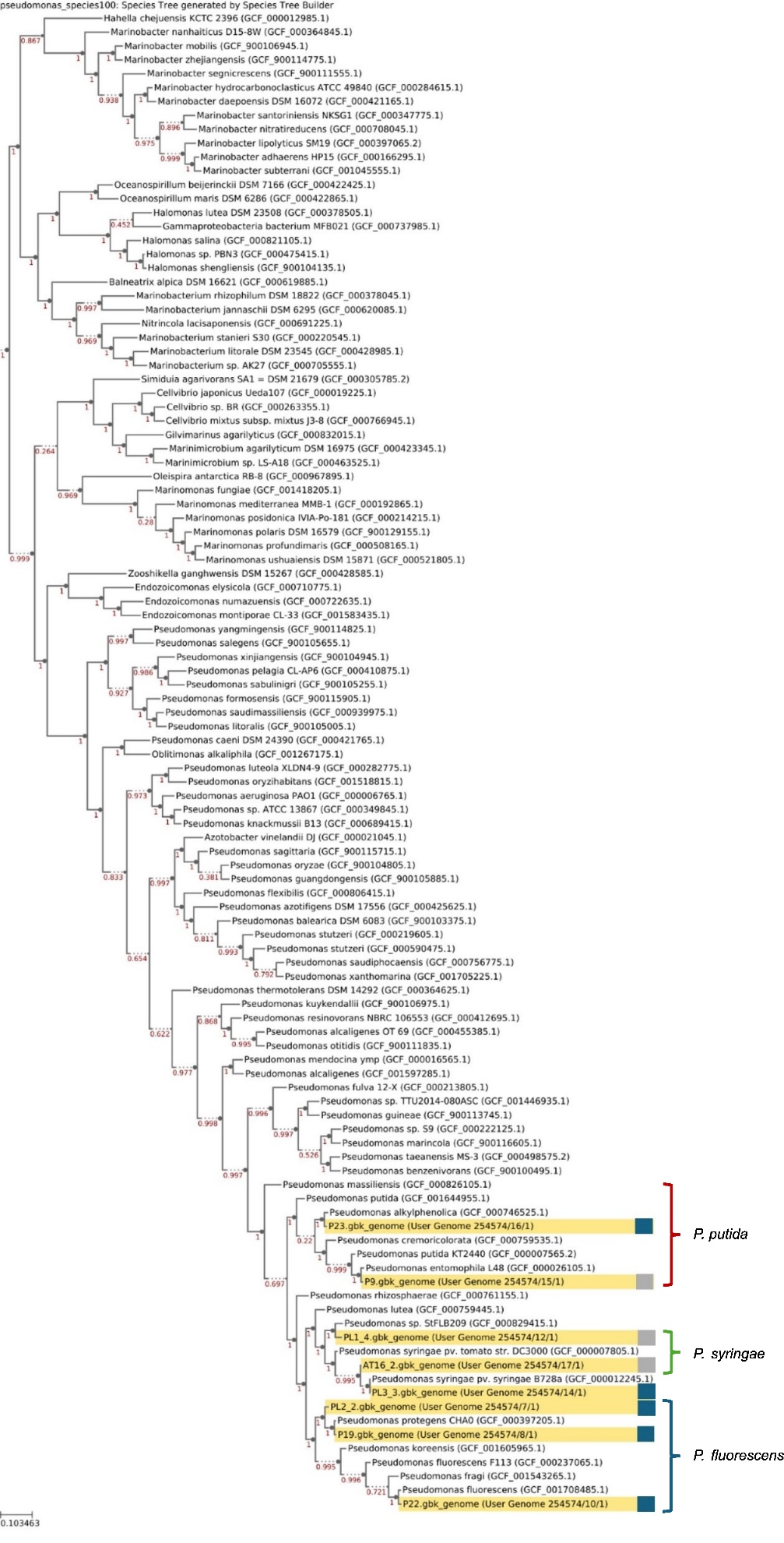

**Figure S9. Whole-genome phylogenetic placement of the eight selected *Pseudomonas* isolates (in yellow) in relation to reference genomes.** Inhibitory strains are indicated by blue and non-inhibitory by grey squares. The species tree is based on 49 concatenated conserved COGs. Branch support values are shown at nodes.

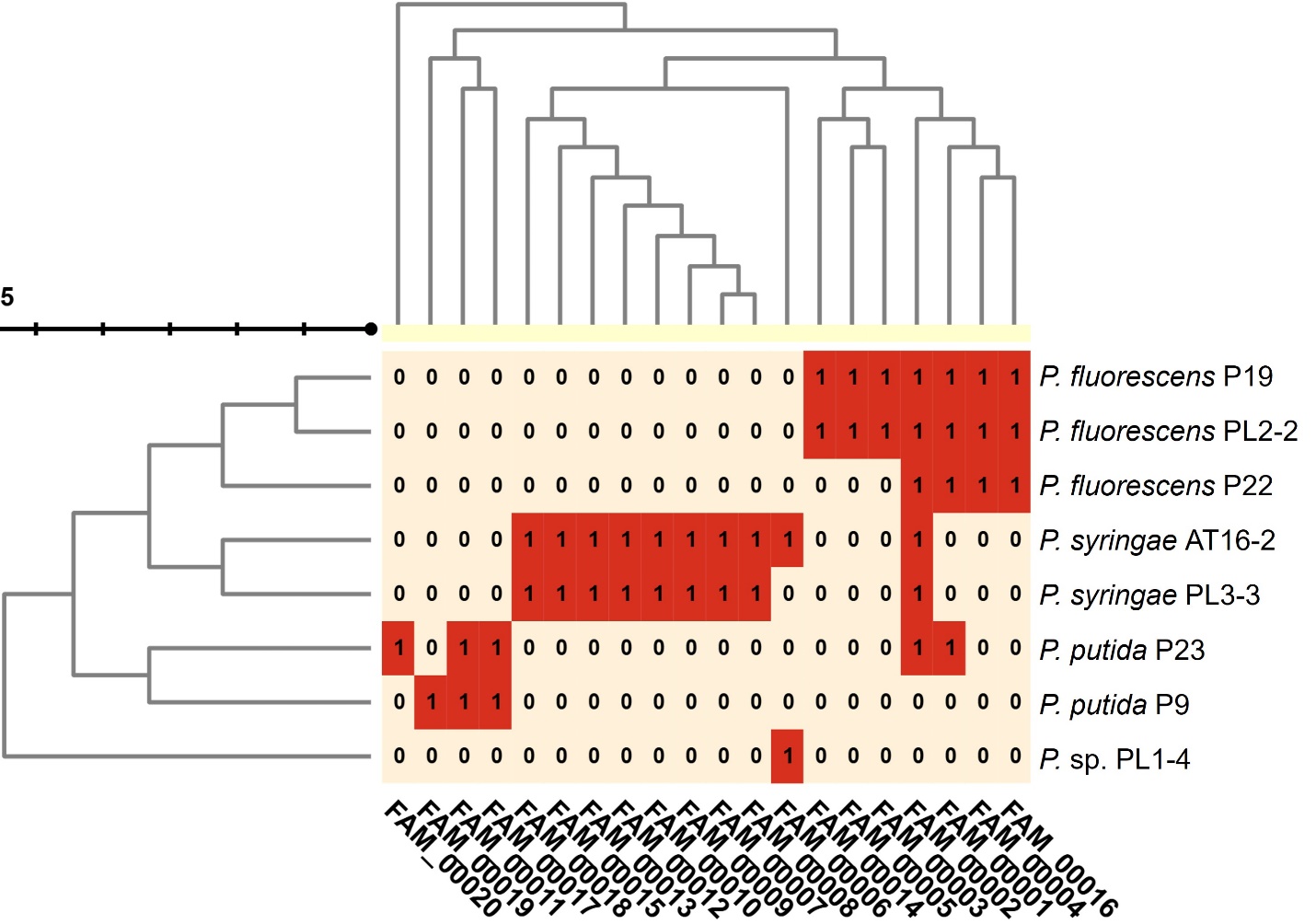

**Figure S10.** **Clustering of selected *Pseudomonas* isolates based on shared biosynthetic gene cluster family (GCF) composition.** The heatmap shows the presence (1, red) or absence (0, light background) of shared GCFs across the eight sequenced isolates.
